# Interpretable adenylation domain specificity prediction using protein language models

**DOI:** 10.1101/2025.01.13.632878

**Authors:** Abhinav K. Adduri, Andrew T. McNutt, Caleb N. Ellington, Krish Suraparaju, Nan Fang, Donghui Yan, Benjamin Krummenacher, Sitong Li, Camilla Bodden, Eric P. Xing, Bahar Behsaz, David Koes, Hosein Mohimani

## Abstract

Natural products have long been a rich source of diverse and clinically effective drug candidates. Non-ribosomal peptides (NRPs), polyketides (PKs), and NRP-PK hybrids are three classes of natural products that display a broad range of bioactivities, including antibiotic, antifungal, anticancer, and immunosuppressant activities. However, discovering these compounds through traditional bioactivity-guided techniques is costly and time-consuming, often resulting in the rediscovery of known molecules. Consequently, genome mining has emerged as a high-throughput strategy to screen hundreds of thousands of microbial genomes to identify their potential to produce novel natural products. Adenylation domains play a key role in the biosynthesis of NRPs and NRP-PKs by recruiting substrates to incrementally build the final structure. We propose MASPR, a machine learning method that leverages protein language models for accurate and interpretable predictions of A-domain substrate specificities. MASPR demonstrates superior accuracy and generalization over existing methods and is capable of predicting substrates not present in its training data, or zero-shot classification. We use MASPR to develop Seq2Hybrid, an efficient algorithm to predict the structure of hybrid NRP-PK natural products from microbial genomes. Using Seq2Hybrid, we propose putative biosynthetic gene clusters for the orphan natural products Octaminomycin A, Dityromycin, SW-163B, and JBIR-39.

## Introduction

More than half of all drugs approved by the Food and Drug Administration (FDA) are derived from bioactive natural products (1, 2). Refined through millions of years of natural selection, bioactive natural products are a valuable source of drug candidates with potentially novel mechanisms of action. For example, non-ribosomal peptides (NRPs) are a class of peptidic natural products that contain many of the drug molecules from the World Health Organization (WHO) list of essential medicines (3). Polyketides (PKs) are another well-studied class of natural products that comprise 20% of the top-selling pharmaceuticals (4) and also display a wide spectrum of bioactivities (5, 6). Despite their structural differences, NRPs and PKs are both assembled via similar mechanisms in bacteria and fungi, enabling the synthesis of diverse NRP-PK hybrid molecules (7, 8), such as the immunosuppressant rapamycin (9) and the anticancer bleomycin (10). Together, NRPs, PKs, and hybrid molecules represent a valuable source of therapeutically relevant drugs (Fig. 1a).

**Fig. 1.**
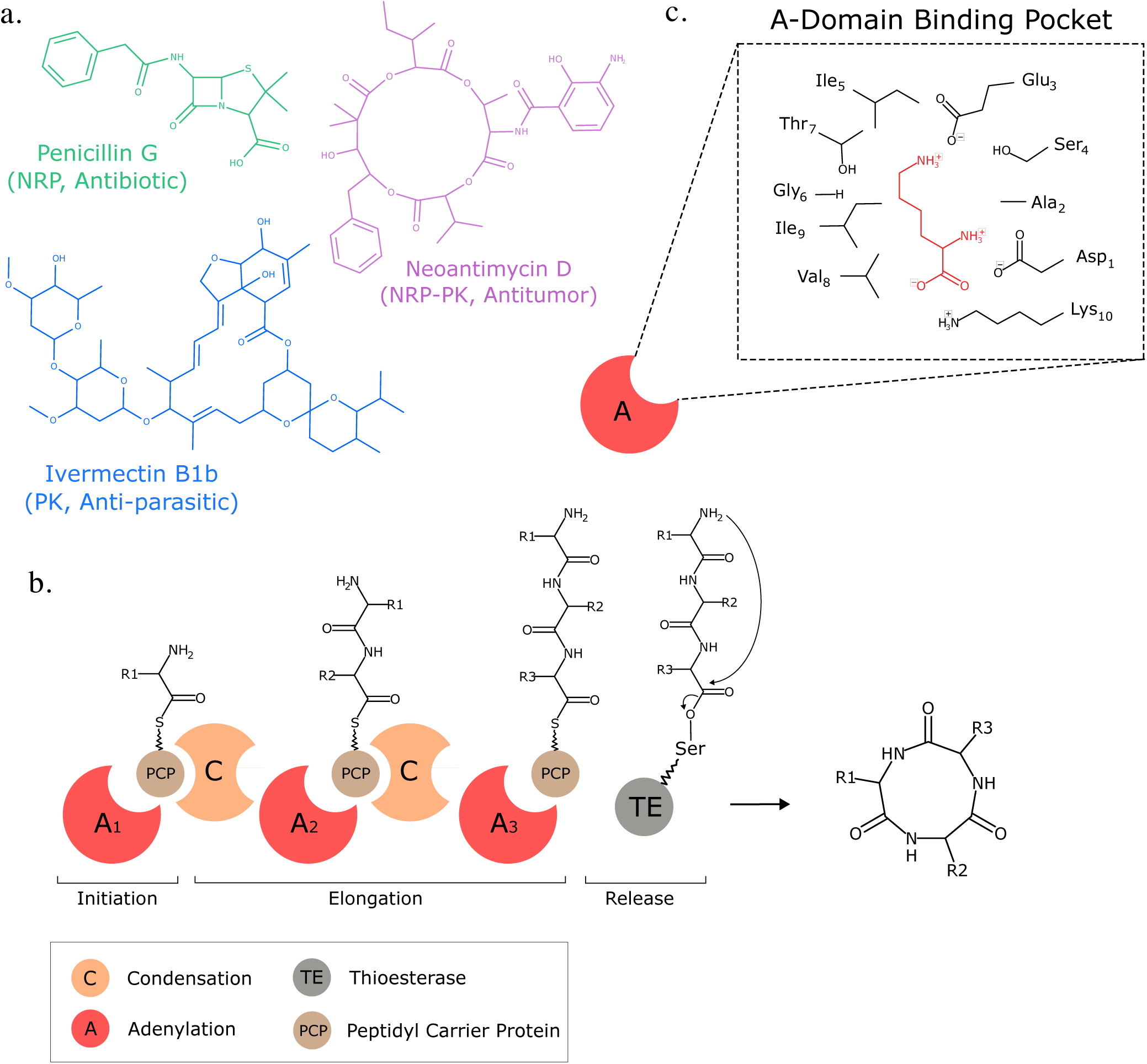
Assembly line enzymology produces diverse and therapeutically valuable natural products. a) NRPs, PKs, and NRP-PK molecules have diverse structures and activities. b) Biosynthetic enzymes act in a coordinated assembly-line fashion to produce NRP-PK hybrids. Adenylation (A-) domains load specific monomers onto PCP domains. Condensation (C-) domains are responsible for linking monomers across adjacent PCP domains to incrementally build the molecule. The process repeats until a thioesterase (TE) domain facilitates the release of the final product. c) A close-up view of the Stachelhaus residues within the binding pocket of an example A-domain. These residues are specificity-conferring and enable different A-domains to recruit different amino acids as needed.

In the past two decades, thousands of NRPs, PKs, and hybrid molecules have been linked to their biosynthetic gene clusters (BGCs), or co-located genes that synthesize natural products (11). However, an analysis of publicly available genome sequencing data revealed hundreds of thousands of BGCs that are not linked to any known compounds (12, 13), representing the enormous potential for novel discovery. Accordingly, several genome mining approaches have been proposed for identifying BGCs (14), predicting the bioactivities of the encoded natural products (15), linking putative BGCs to known natural products (16), and predicting the structure of natural products encoded by putative BGCs (17–19). As paired tandem mass spectrometry and genome sequencing data are now readily obtainable for microbial isolates and communities (20, 21), recent methods adopt a multi-omics approach by predicting a large set of putative natural products for a given BGC and filtering the predictions using paired tandem mass spectrometry data (22, 23). Despite these advances, a survey of current literature revealed that existing genome mining tools are significantly better at mining NRPs or PKs from BGCs than NRP-PK hybrid molecules (24), which remain underrepresented and difficult to predict due to their structural and biosynthetic complexity (7, 25–27).

NRPs, PKs, and their hybrids are synthesized through the coordinated action of enzymes arranged in an assembly-line fashion within BGCs (Fig. 1b). In NRPs, adenylation (A-) domains within the BGC are responsible for incrementally adding specific amino acids or hydroxy acids to a growing peptide (26). Analogously in PKs, acyltransferase (AT-) domains sequentially add specific alpha-carboxyacyl (ketide) subunits to the final structure (26, 28). Therefore, computational approaches for predicting NRP, PK, or NRP-PK hybrid structures encoded by a given BGC are limited by the accuracy of A-domain and AT-domain substrate specificity prediction from their amino acid sequences. Yadav et al. identified a 24 amino acid motif of specificity-conferring residues within the AT-domain binding pocket and achieved an impressive 95% accuracy in substrate prediction (29).

Similarly, for A-domains, Stachelhaus et al. reported a 10 amino acid specificity-conferring motif, or “Stachelhaus code”, in the A-domain binding pocket (Fig. 1c), which achieved 86% accuracy in classifying substrate specificity for 160 A-domains (30). As the amount of training data for A-domain binding specificity increased, researchers observed an increasing number of A-domains that share identical Stachelhaus codes yet display different substrate specificities (31).

To better differentiate between A-domains with identical Stachelhaus codes, Rottig et al. expanded the code to 34 residues within 8Å of the binding pocket to capture the context around the Stachelhaus code (32), enabling machine learning methods to predict the specificity based on the 8Å signature (32, 33). However, while this approach improved prediction accuracy for some A-domains, it did not fully resolve the challenges posed by inherently promiscuous A-domains, which can recruit multiple substrates despite having identical amino acid sequences (34, 35).

Furthermore, later work revealed that methods for A-domain specificity prediction were severely overfitted to their training data, with validation accuracy on out-of-distribution test data as low as 22%, resulting in poor overall performance on novel BGCs (35). This suggests that A-domain specificity prediction is a weak link in novel NRP and NRP-PK structure prediction (22). Our results show that A-domain specificity prediction is especially poor for NRP-PK hybrids, which can incorporate rare, non-standard amino acids.

To more systematically capture the context of amino acids in the binding pocket, in this work we explore the use of protein language models to featurize the A-domain. Protein language models have proven to be effective foundation models in biology, as they learn characteristics of amino acid sequences over millions of protein sequences (36–39). Despite learning these characteristics in an unsupervised fashion with no structural information, these models can capture dependencies between amino acids that are close in three-dimensional space but far apart in sequence space (40). As such, protein language models have been used to guide protein design and generation (41–43) and predict drug-target interactions (44–46).

We propose MASPR (**m**odeling **A**-domain **s**pecificity using unsupervised **p**retrained **r**epresentations), which leverages a protein language model to generate embeddings for A-domains. Building on recent deep learning methods that predict molecular fingerprints (47–49), MASPR employs two neural networks for interpretable A-domain specificity prediction. The first neural network is trained to generate an interpretable molecular fingerprint of the substrate recruited by a given A-domain. To accommodate promiscuous A-domains that may interact with multiple substrates, a second neural network is trained on the predicted fingerprints from the first neural network and the target fingerprints to learn a latent substrate embedding that represents potential A-domain binding partners. The latent substrate embedding is a data-driven, compact representation learned by the model that encodes the most likely substrates for a given A-domain.

MASPR predicts specificity via nearest substrate search through a precomputed database of these latent substrate embeddings. MASPR is further trained to compute latent embeddings for substrates not present in the training data, meaning this database can include novel substrates as specified by the user. This enables MASPR to perform interpretable predictions of substrate specificities not found in the training data, or zero-shot classification. MASPR achieves state-of-the-art accuracy, improving top-5 accuracy from 47.5% to 63.1% in out-of-distribution generalization and from 67.8% to 72.2% on promiscuous A-domains. In a leave-one-substrate-out cross-validation designed to measure zero-shot predictive performance, MASPR achieved over 50% top-5 accuracy for more than half of the held-out substrates.

We then used MASPR to develop Seq2Hybrid, a genome mining method for predicting mature modular type 1 NRP-PK hybrid structures directly from microbial genomes. Seq2Hybrid uses MASPR to annotate A-domain specificities in hybrid BGCs and accounts for biosynthetic uncertainties such as A-domain promiscuities, variable gene assembly orders, and post-assembly enzymatic modifications by outputting a database of potential encoded natural products. Seq2Hybrid can subsequently filter the database to retain only molecules with sufficient spectral evidence if paired mass spectrometry (MS) data is available. We demonstrate that even in the absence of paired MS data, Seq2Hybrid with MASPR outperforms existing methods at recovering encoded natural products. Together, MASPR and Seq2Hybrid enable state-of-the-art A-domain specificity prediction and genome mining for NRP-PK hybrid molecules encoded by microbial BGCs.

## Results

### Overview of MASPR algorithm

MASPR utilizes protein language models for interpretable and accurate prediction of A-domain specificities (Fig. 2). Rather than performing classification over a fixed set of substrates, MASPR converts substrates to their fingerprint representations, which are used as regression targets during training. The molecular fingerprint used for a given substrate is a concatenation of the MACCS key (50), the ECFP4 fingerprint calculated as the Morgan fingerprint with length 128 and radius 2 in RDKit (51, 52), and the average partial charge of atoms in the substrate (Fig. 2a). For a given A-domain sequence with length *n*, MASPR inputs the amino acid sequence to an ESM-2 language model (53) to generate an *n ×* 1280 dimensional representation. MASPR extracts the embeddings for the Stachelhaus residues to obtain a 10 *×* 1280 dimensional representation for each A-domain sequence.

**Fig. 2.**
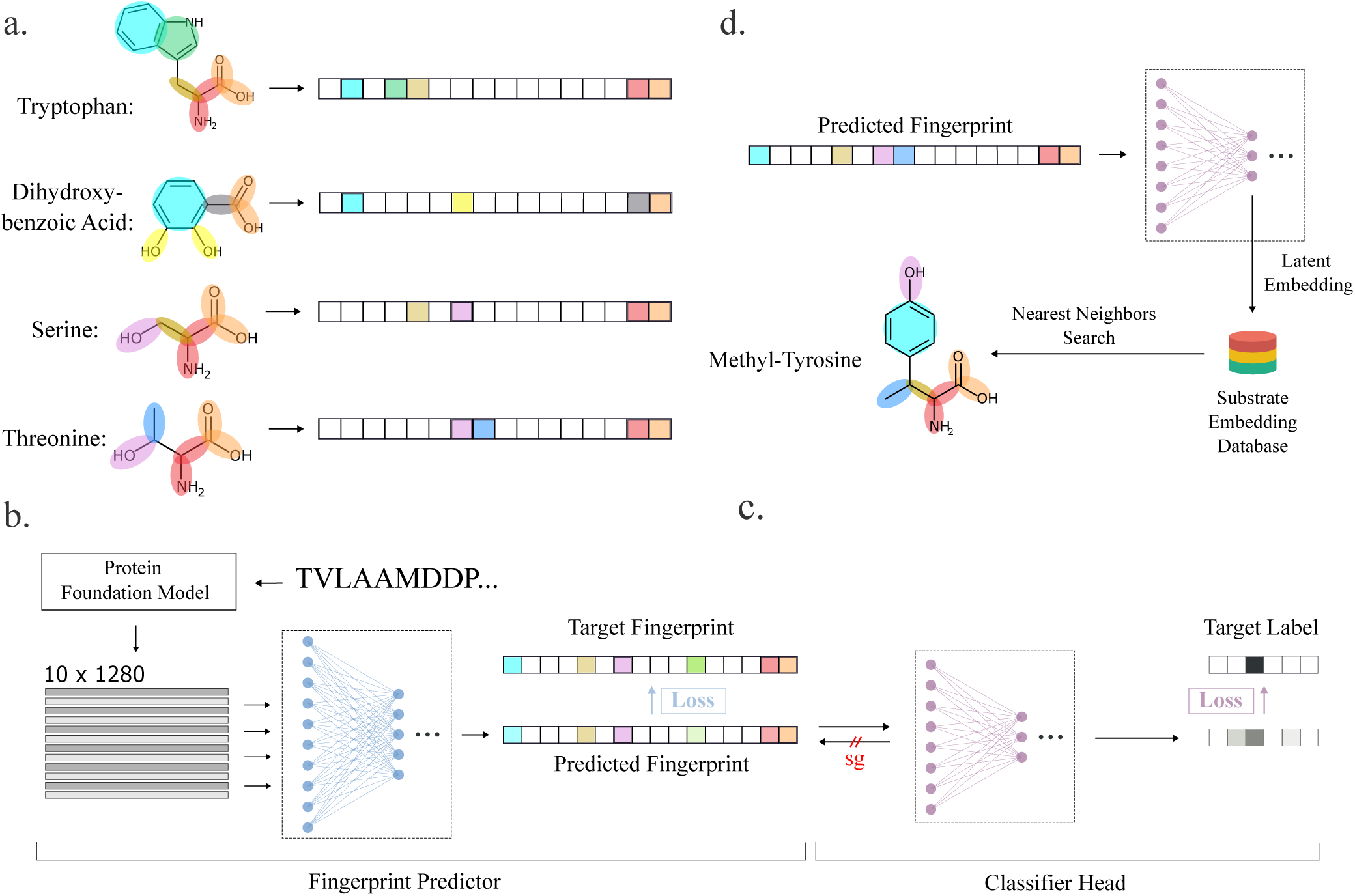
Overview of MASPR. a) Using RDKit, substrates are converted from their SMILES representation to a substructure-based fingerprint, and are augmented with contextual connectivity information by concatenating an ECFP (Morgan) fingerprint. b) An A-domain sequence is inputted to a protein language model (PLM) to obtain embeddings for the Stachelhaus residues, resulting in a 10 *×* 1280 dimensional representation for each input sequence. These representations are used as inputs to the first neural network (fingerprint predictor), which is trained to predict the substructure-based fingerprint. c) Since nearest neighbor search in fingerprint space cannot account for promiscuous A-domains, which may recruit substrates with dissimilar fingerprints, a second neural network (classifier head) is trained to recover the substrate labels from the predicted fingerprints and target fingerprints. The classifier head may also be trained on fingerprints for substrates not found in the training data, as these gradients do not flow back through the fingerprint predictor (stop-gradient, or sg, in the figure). d) At test time, the hidden representation of the classifier head is used for nearest neighbor search for classification, enabling zero-shot classification of substrates not in the training data. The fingerprint predictor neural network output is used to highlight substructural features relevant to the final prediction.

MASPR trains the first neural network (fingerprint predictor) to recover molecular fingerprints by minimizing the cosine distance between the predicted and actual fingerprints (Fig. 2b). Then, MASPR trains a second neural network (classifier head) to predict the substrate labels from the predicted fingerprints (Fig. 2c). Because the gradients from the classifier head do not flow back to the fingerprint predictor, the classifier head can also be trained on the target fingerprints, as well as chemical fingerprints for substrates not found in the training data.

At test time, MASPR computes the latent embedding for an input A-domain and retrieves the top-*k* nearest substrates from an embedding database, where substrate distance is calculated as the cosine distance of their embeddings (Fig. 2d). MASPR can compute embeddings for substrates not present in the training data using their chemical fingerprints, enabling zero-shot prediction of novel substrates. MASPR can additionally use the predicted molecular fingerprint to identify the substructural features that were most relevant in its predictions.

### MASPR outperforms existing methods in A-domain specificity prediction

Previous work on A-domain substrate prediction accuracy showed that accuracy is often overestimated due to A-domains in the test set that are very similar to A-domains in the training set (35). Therefore, we stratify the test set into buckets, where a test A-domain is in bucket *B_i_*_+_ if its 8Å signature has a Hamming distance of at least *i* residues from any 8Å signature in the training set, where the Hamming distance measures the number of positions at which two 8Å signatures differ. MASPR was benchmarked using ESM-2 and AlphaFold2 (AF2) featurizations of the Stachelhaus residues (36, 54) and a one-hot encoding of the 8Å signature (Fig. 3). AlphaFold2 (AF2) features were obtained using ColabFold (55) for input A-domain sequences by extracting the hidden layer single representation before the AF2 structure modules (54), resulting in 256-dimensional embeddings per residue. PDB entries for A-domains in their adenylating conformations (1AMU, 4D57, 4D56, 3VNS, 3DHV, 4ZXI, 5N9X) were used as templates for ColabFold. ESM-2 features were extracted using the `esm2_t33_650M_UR50D` model, which provides 1280-dimensional embeddings per residue without templates.

**Fig. 3.**
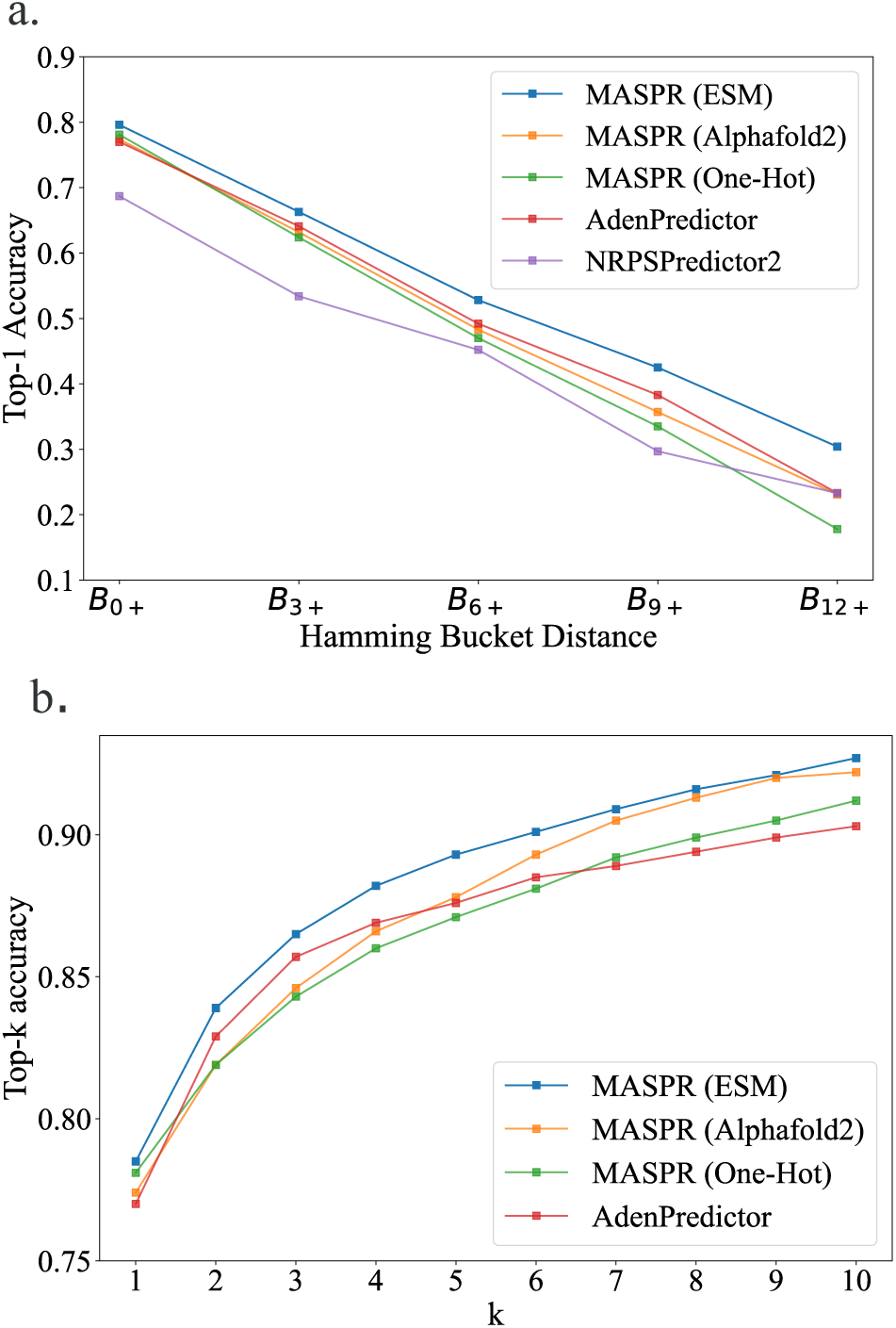
Accuracy of MASPR vs other methods. a) Top-1 accuracy across different Hamming buckets, representing increasing dissimilarity to the training data. Bucket *B_i_*_+_ corresponds to the portion of test data with Hamming distance of *i* or higher from all training data points, where the Hamming distance measures the number of positions at which two 8Å signatures differ. On average across all train/test splits, the Hamming buckets represent 459, 191, 105, 64, and 30 data points, respectively. b) Top-*k* accuracy for different values of *k*. MASPR with ESM-2 featurization is the best-performing method for all values of *k*.

Accuracies are reported after averaging across 12 splits of the training and test data (Fig. 3). MASPR with ESM-2 featurization outperforms AdenPredictor, the previous state-of-the-art, across all test buckets (Fig. 3a), and achieves higher top-*k* accuracy for all values of *k* (Fig. 3b) despite the relatively small amount of training data (2294 training data points). Furthermore, the performance gap between MASPR and AdenPredictor widens as the bucket distance increases, with MASPR outperforming by up to 7% on test points in *B*_12+_, demonstrating that MASPR can generalize better to out-of-distribution test data. Despite using the same architecture, MASPR performance drops significantly when using a one-hot encoding (where each amino acid is represented by a binary vector with a 1 in a unique position and 0 elsewhere) of the 8Å signature. This demonstrates that the ESM-2 featurization contains a signal relevant to A-domain substrate specificity and enables high sample efficiency. We also tested larger ESM models which output higher dimensional embeddings per residue, but noticed worse overall performance, possibly due to the scarcity of training data relative to the embedding size.

### MASPR improves generalization and accuracy

One drawback of the neural network for predicting fingerprints is its tendency to generate averaged fingerprints for promiscuous A-domains that recruit diverse substrates with dissimilar fingerprints. At test time, this leads to reduced accuracy when performing the nearest substrate search in fingerprint space. MASPR addresses this by training a second neural network, the classifier head, that predicts substrate labels from the predicted fingerprints and the correct fingerprints (computed by RDKit). While the output of the classifier head is discarded after training, the learned hidden representation serves as a substrate embedding whose metric properties are more suitable for representing A-domain specificity and promiscuity. Indeed, removing the classifier head and directly using the molecular fingerprint for the nearest substrate search leads to a significant drop in top-*k* accuracy (Fig. 4a). To further evaluate the impact of the classifier head for promiscuous A-domains, we stratified the dataset by training on all non-promiscuous sequences. Then, for each promiscuous A-domain sequence with observed specificity for *n* substrates, we randomly selected one sequence-substrate pair to add to the training set and used the remaining *n −* 1 sequence-substrate pairs for the test set. Our results demonstrate that MASPR with the classifier head consistently outperforms both MASPR without the classifier head and AdenPredictor across all Hamming bucket distances and top-k accuracy metrics (Fig. 8).

**Fig. 4.**
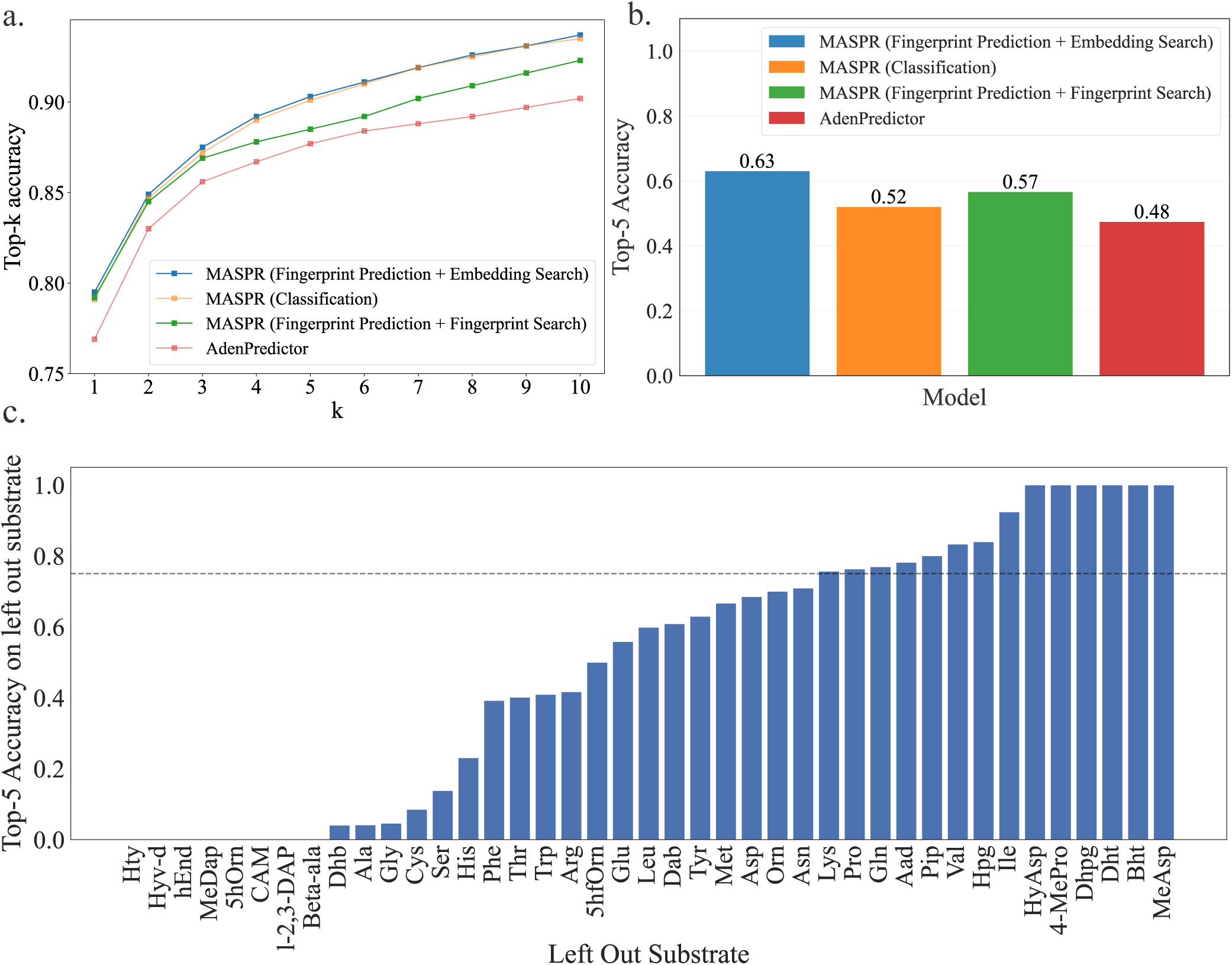
Regression and classification objectives synergistically improve MASPR generalization and accuracy. a) MASPR models that use predicted molecular fingerprints for nearest substrate search have poor top-*k* performance compared to models that use latent embeddings from the classifier head. b) When trained on bacterial A-domain data and tested on fungal data, MASPR models that predict fingerprints can generalize better than models that do not. c) In a leave-one-substrate-out cross-validation, MASPR with ESM-2 achieves over a top-5 prediction accuracy of at least 75% for 34% of substrates.

**Fig. 5.**
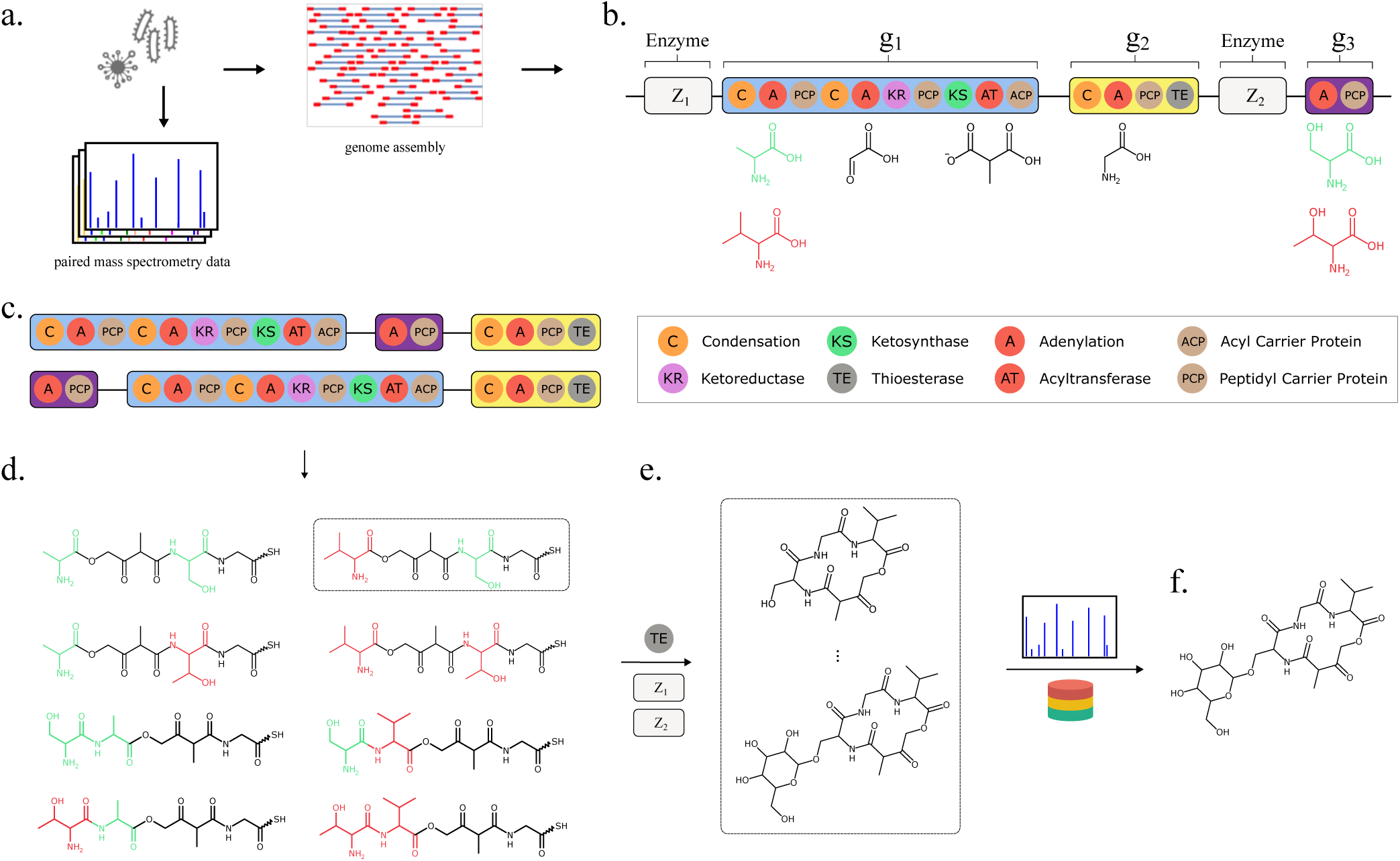
Overview of Seq2Hybrid. a) Genomic DNA and paired mass spectrometry data are collected from microbial strains. b) Given a microbial genome as input, Seq2Hybrid searches for NRP-PK hybrid BGCs and for enzymes that perform post-assembly modifications. Each A-domain is annotated with the most likely set of monomers it will incorporate using MASPR. AT-domains are annotated using Minowa et al. (17). c) Different assembly orders are calculated from the BGC. d) For each assembly order and each monomer assignment for an active domain, Seq2Hybrid generates core molecules. e) The core molecules are modified post-assembly by enzymes in the BGC to generate a database of hypothetical natural products. f) Hypothetical natural products are further searched against mass spectra, if provided, and high-scoring matches are retained (56).

To explore the role of fingerprint prediction and the classifier head on accuracy and generalization, we train models on bacterial A-domain sequences and test them on fungal A-domain sequences, under the hypothesis that a model that captures true binding dynamics of A-domains should be able to generalize despite evolutionary differences. MASPR achieved 15% higher top-5 accuracy than AdenPredictor in this benchmark. MASPR models that integrate fingerprint prediction with a classifier head for latent space nearest substrate search outperformed models that solely relied on nearest fingerprint search, as well as models that replaced nearest neighbor search with direct classification over a fixed set of substrates (Fig. 4b). Our results suggest that predicting fingerprints and using the learned latent space of the classifier head for the nearest substrate search synergistically enhance generalization.

### MASPR classifies unseen substrate specificities

Because the classifier head is trained on fingerprints (generated by RDKit from SMILES representations), MASPR can compute embeddings for substrates not included in the training data from their SMILES representations, enabling zero-shot prediction of novel substrates. To evaluate MASPR’s zero-shot predictive accuracy, we use a leave-one-substrate-out strategy, in which the model is trained on all substrate labels except one and tested solely on A-domains that recruit the omitted substrate label. MASPR achieves a top-5 prediction accuracy of at least 75% for 34% of substrates, and a top-5 prediction accuracy of at least 50% for over half of the left-out substrates (Fig. 4c). None of the other methods have the capacity for zero-shot predictions.

### Incorporating knowledge about binding-pocket residues enhances predictive accuracy

Previous methods have used averaging to combine per-residue features across the whole protein for binding prediction (44). We observe that MASPR performance drops significantly when averaging across the whole protein (Fig. 9). Interestingly, averaging across only the Stachelhaus residues recovers much of the performance lost by whole protein averaging, which suggests that the embeddings for individual Stachelhaus residues carry signals relevant to A-domain specificity. Although previous approaches exclude the tenth Stachelhaus residue due to its invariant Lysine identity, including it led to slightly better performance in our experiments, likely due to the context-dependent nature of protein language model embeddings. Maintaining a separate channel for each Stachelhaus residue (10 *×* 1280) results in the best performance, suggesting that, when possible, the incorporation of known binding pocket information can significantly improve substrate specificity prediction, especially when the size of the training data is small.

### MASPR enables more accurate NRP-PK structure predictions

Seq2Hybrid is an end-to-end tool that leverages MASPR for the prediction of mature NRP-PK hybrid molecules (Fig. 5). Starting with a microbial genome as input (Fig. 5a), Seq2Hybrid searches for BGCs in the genome that potentially encode NRP-PK hybrids (Fig. 5b). These are identified as BGCs that contain active domains (A-domains or AT-domains), which recruit monomers into the natural product. In the case of NRP-PK hybrids, these monomers are usually either amino acids, hydroxy acids, or *α*-carboxyacyl-CoA extender units (ketides). Seq2Hybrid uses MASPR to predict the top three most likely monomers that each A-domain might add and uses existing approaches (17) to predict the most likely monomer that each AT-domain might add (Fig. 5b). Then, Seq2Hybrid computes biosynthetic assembly lines, which are defined as a particular ordering of biosynthetic genes in the product assembly (Fig. 5c). For each assembly line, Seq2Hybrid uses the predicted active domain specificity to produce a list of precursor hybrid molecules (Fig. 5d). Finally, for each precursor hybrid molecule, Seq2Hybrid combinatorically applies various post-assembly modifications to generate a database of mature NRP-PK hybrid predictions (Fig. 5e). If paired mass spectrometry data is also provided, Seq2Hybrid further searches NRP-PK hybrid predictions against mass spectra and retains the high-scoring matches (Fig. 5f).

### Benchmarking Seq2Hybrid

Seq2Hybrid was bench-marked on 286 NRP-PK hybrid molecules in MIBiG (11) for which PRISM 4 (24) and antiSMASH 7.0 (14) structural predictions were available using only the genome mining module (no paired mass spectrometry data was provided for fair comparison). We further focused only on molecules with a type-1 polyketide component. To ensure no leakage between train and test sets in our data collection for tailoring modifications, we used 65 hybrid BGCs added in MIBiG 3.0 that were not present in MIBiG 2.0 as test BGCs to measure the out-of-distribution performance of each method. These BGCs are also not present in the PRISM 4 training data.

For each BGC, the best Seq2Hybrid prediction was compared to the best PRISM prediction and the best anti-SMASH prediction, where the best prediction for a given method was computed using Tanimoto similarity against the ground truth NRP-PK. Tanimoto similarity was calculated using Morgan fingerprints with 1024 bits and a radius of 3. For each method, we calculated the number of hybrid BGCs for which the Tanimoto similarity of the ground truth and the best-predicted molecule was at or above a given threshold. Seq2Hybrid outperforms PRISM and antiSMASH across all Tanimoto thresholds (Fig. 6). At a Tanimoto similarity threshold of 0.7, Seq2Hybrid identifies 25 molecules, while PRISM 4 identifies two molecules and antiSMASH does not identify any molecules (Supplementary Table S1). Though the performance of both methods suffers on the test set, which contains several unseen chemical modifications, Seq2Hybrid maintains a similar relative performance improvement over PRISM 4.

**Fig. 6.**
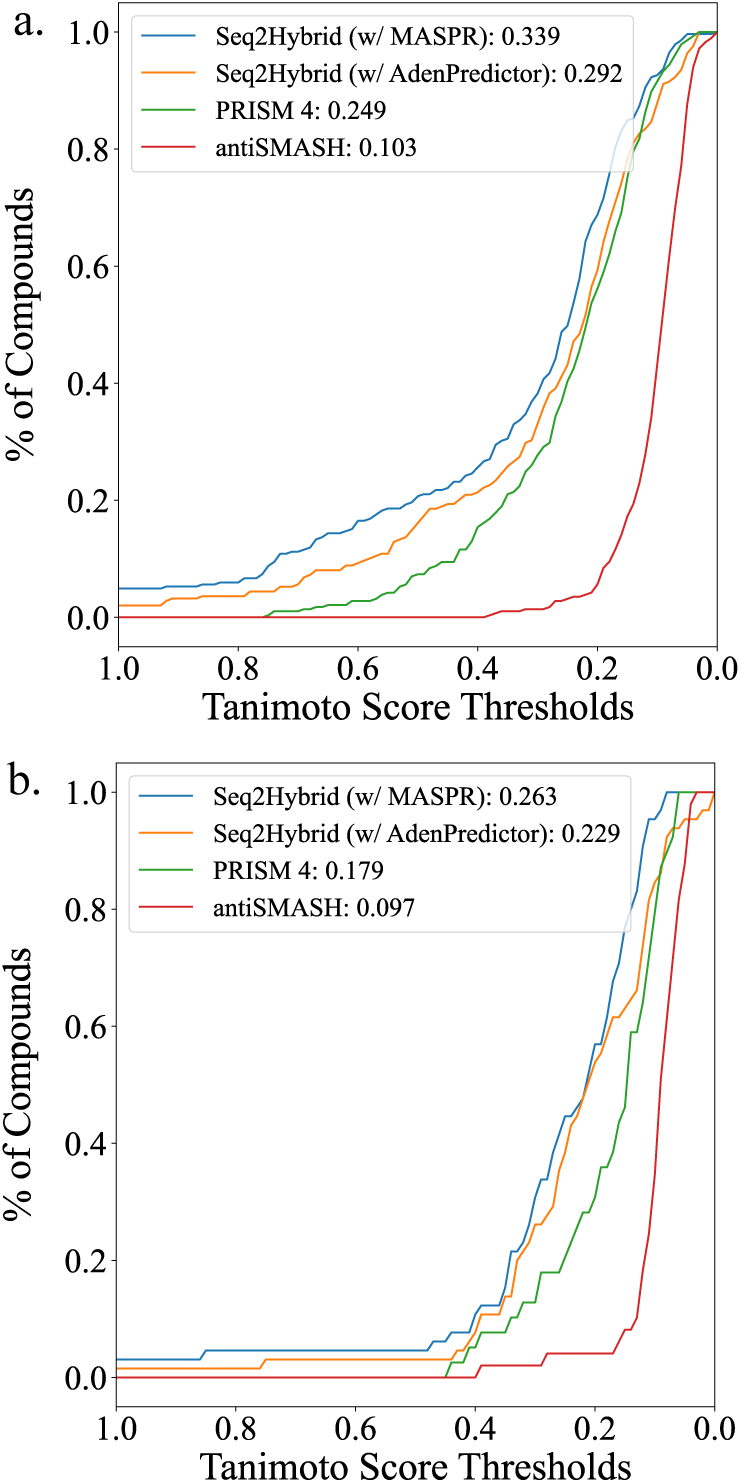
Tanimoto Comparison of Seq2Hybrid, PRISM and anti-SMASH. Seq2Hybrid can accurately recover more NRP-PK hybrids than PRISM across all measured Tanimoto thresholds, and using Seq2Hybrid with MASPR further improves performance over using AdenPredictor (35). AntiSMASH only reports core structure but is included as a baseline to show the importance of accounting for tailoring modifications. The Tanimoto similarity of the best prediction for each method against the ground truth is reported in the legend, averaged across all BGCs in a) the training set, and b) the test set.

**Fig. 7.**
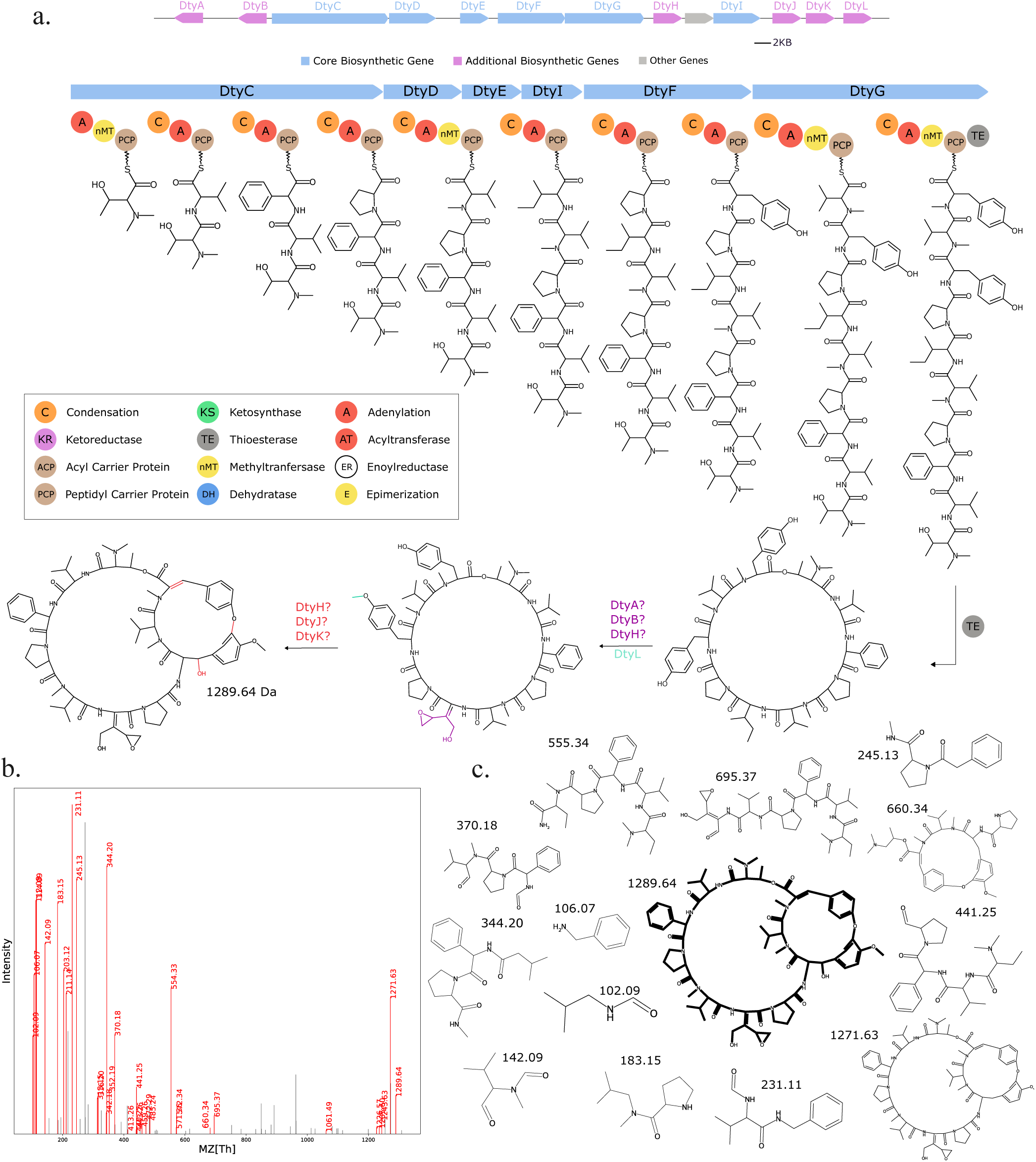
Seq2Hybrid predicts Dityromycin BGC. a) The predicted biosynthetic pathway for Dityromycin. *DtyH*, *DtyJ*, and *DtyK* are all cytochrome p450 enzymes, and are likely responsible for hydroxylation and cross-linkage of the Tyrocine residues. *DtyA* and *DtyB* are 2-oxo-acid dehydrogenases and enoyl-CoA hydratases, respectively, and in conjunction with the p450 enzymes, are likely responsible for the modification of Isoleucine to E-2-amino-3-hydroxymethyl-4,5-epoxy-*α*,*β*-dehydropentanoic acid. *DtyL* is a methyltransferase that methylates one of the Tyrosine residues. b) Paired mass spectrometry data for this molecule was obtained from *Streptomyces kasugaensis* NBRC 13851. c) Annotated mass fragments providing evidence that this molecule, or an isomer, is present in the biological sample.

**Fig. 8.**
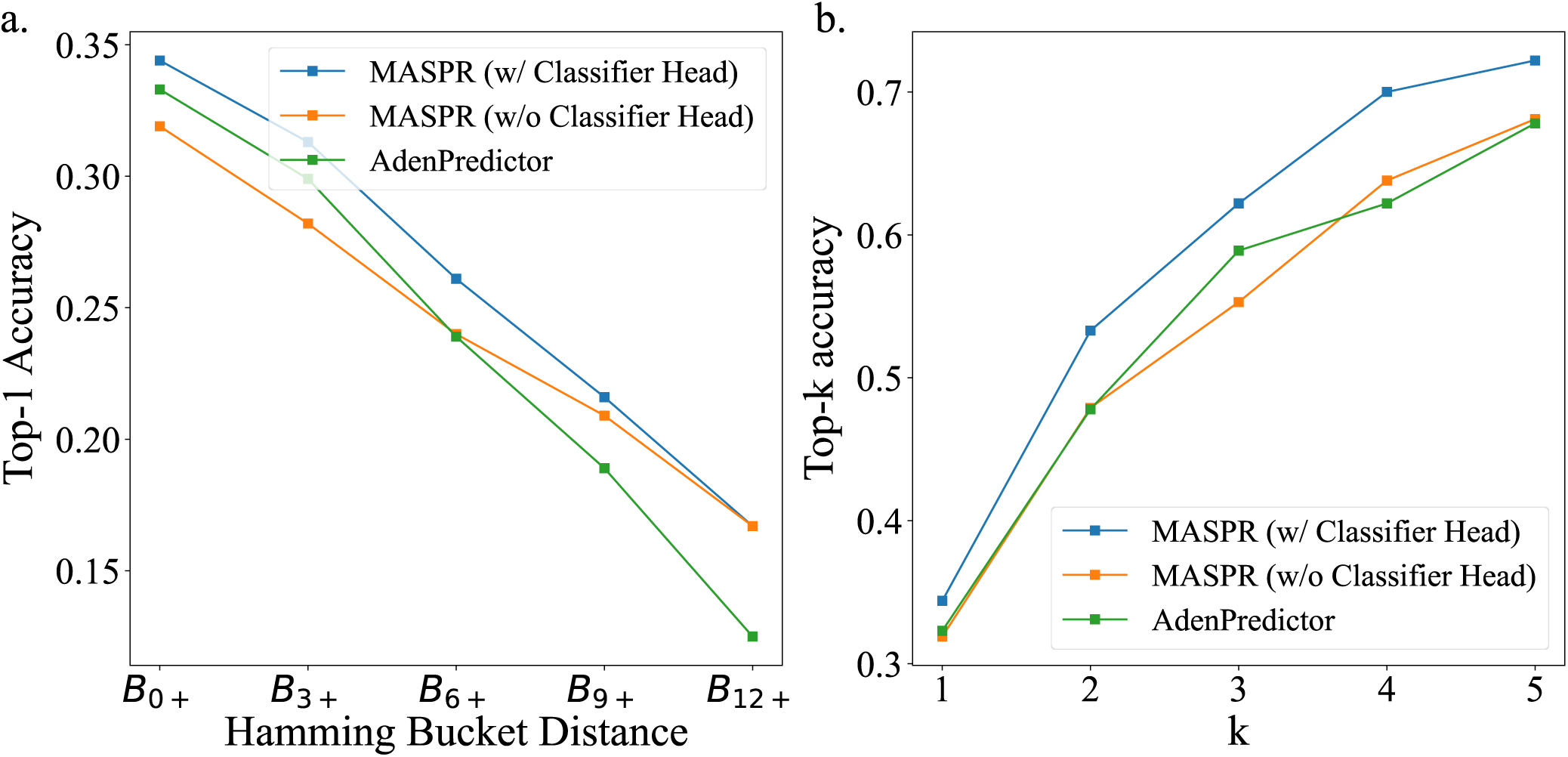
MASPR with a classifier head offers improved accuracy on promiscuous A-domains. a) Top-1 accuracy across Hamming bucket distances for MASPR (with and without classifier head) and AdenPredictor. Higher Hamming bucket distances indicate greater dissimilarity from training data. b) Top-k accuracy for k=1 to 5, comparing the performance of MASPR variants and AdenPredictor. In both metrics, MASPR with the classifier head consistently outperforms the other methods.

**Fig. 9.**
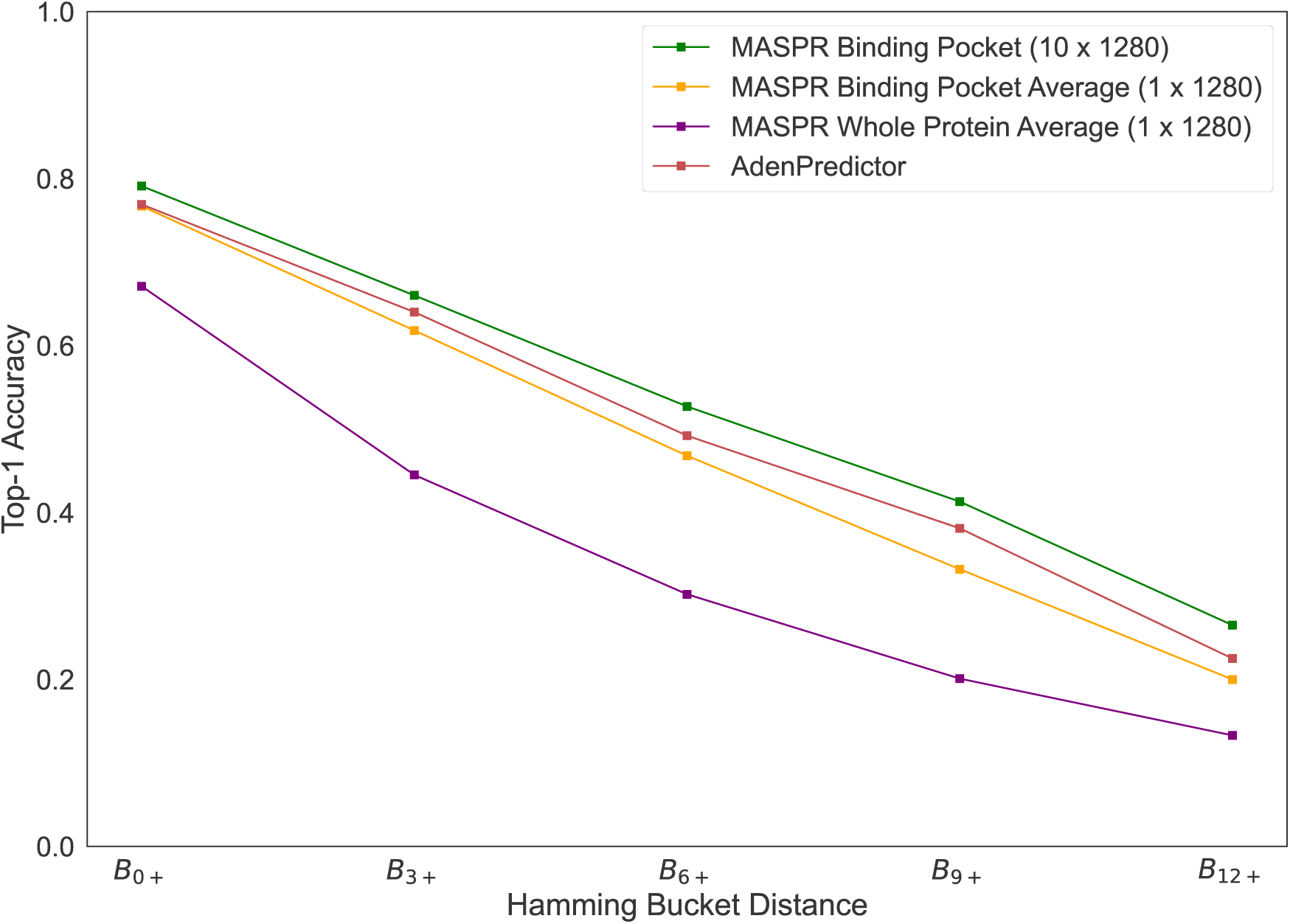
Accuracy of MASPR with different featurizations. Averaging the ESM embeddings across all residues in the proteins results in significantly worse predictive accuracy. Averaging across only the Stachelhaus residues recovers most of the lost accuracy, but still falls short of the accuracy achieved by maintaining a separate channel for each Stachelhaus residue. On average across all train/test splits, the Hamming buckets *B*_0+_, *B*_3+_, *B*_6+_, *B*_9+_, and *B*_12+_ represent 459, 191, 105, 64, and 30 data points, respectively.

It should be emphasized that the main contribution of antiSMASH is genome mining and core structure prediction; therefore it is unfair to compare its performance to Seq2Hybrid and PRISM at predicting mature hybrid compounds. Nevertheless, it is included as a baseline to illustrate the importance of accounting for assembly order and modifications in predicting mature natural product structures. MASPR enables Seq2Hybrid to make predictions with high Tanimoto similarity to the ground truth even when BGCs contain A-domains that recruit substrates not present in the training data, such as 2-amino-6-hydroxy-4-methyl-8-oxodecanoic acid in Leucinostatin (Fig. 10).

**Fig. 10.**
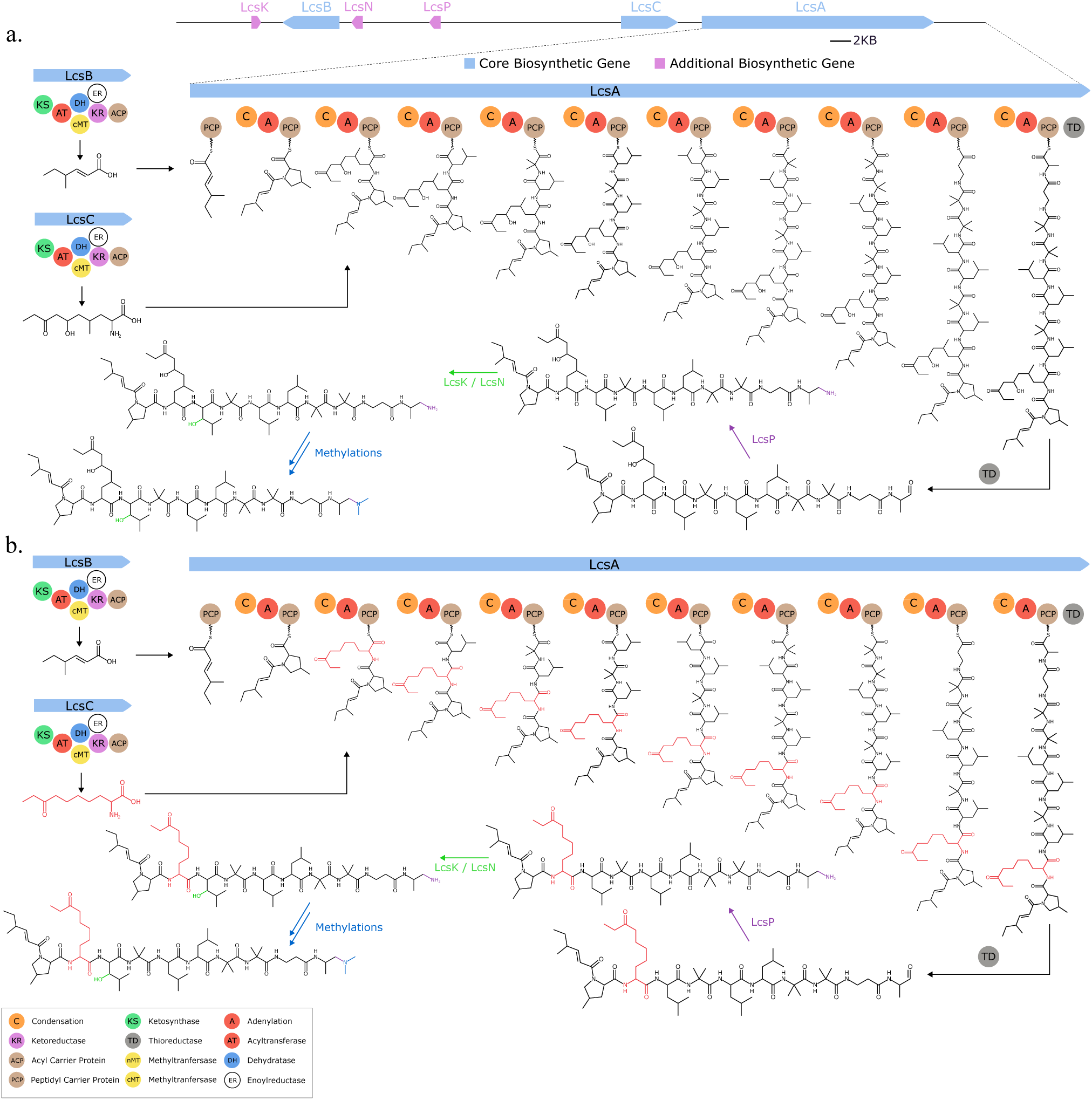
MASPR enables Seq2Hybrid to recover NRP-PK hybrids with rare amino acids. a) The previously reported biosynthetic pathway for Leucinostatin A, which includes *2-amino-6-hydroxy-4-methyl-8-oxodecanoic acid*. b) The biosynthetic pathway assigned by Seq2Hybrid. Although MASPR mispredicts the monomer added by the second A-domain (shown in red), the misprediction is close enough to the ground truth that the final structure still bears high Tanimoto similarity to the ground truth (0.802).

### Seq2Hybrid identifies known hybrid molecules

We used Seq2Hybrid to search mass spectra of eight *Streptomyces* strains against the molecules predicted from their genomes (Supplementary Table S2). Seq2Hybrid correctly identified the structure of known hybrids ilamycin G and rufomycin NBZ8 from *Streptomyces atratus* NBRC 3897 (Fig. 11), pyridomycin from *Streptomyces pyridomyceticus* NRRL B-2517 (Fig. 12), neoantimycin from *Streptomyces orinoci* NBRC 13466 (Fig. 13), and rakicidin B from *Micromonospora chalcea* NRRL B-2672 (Fig. 14) at a Tanimoto threshold of 1.0. Seq2Hybrid also identified a putative BGC for lydiamycin A (57) in *Streptomyces alboflavus* strain MDJK44 (Fig. 15).

**Fig. 11.**
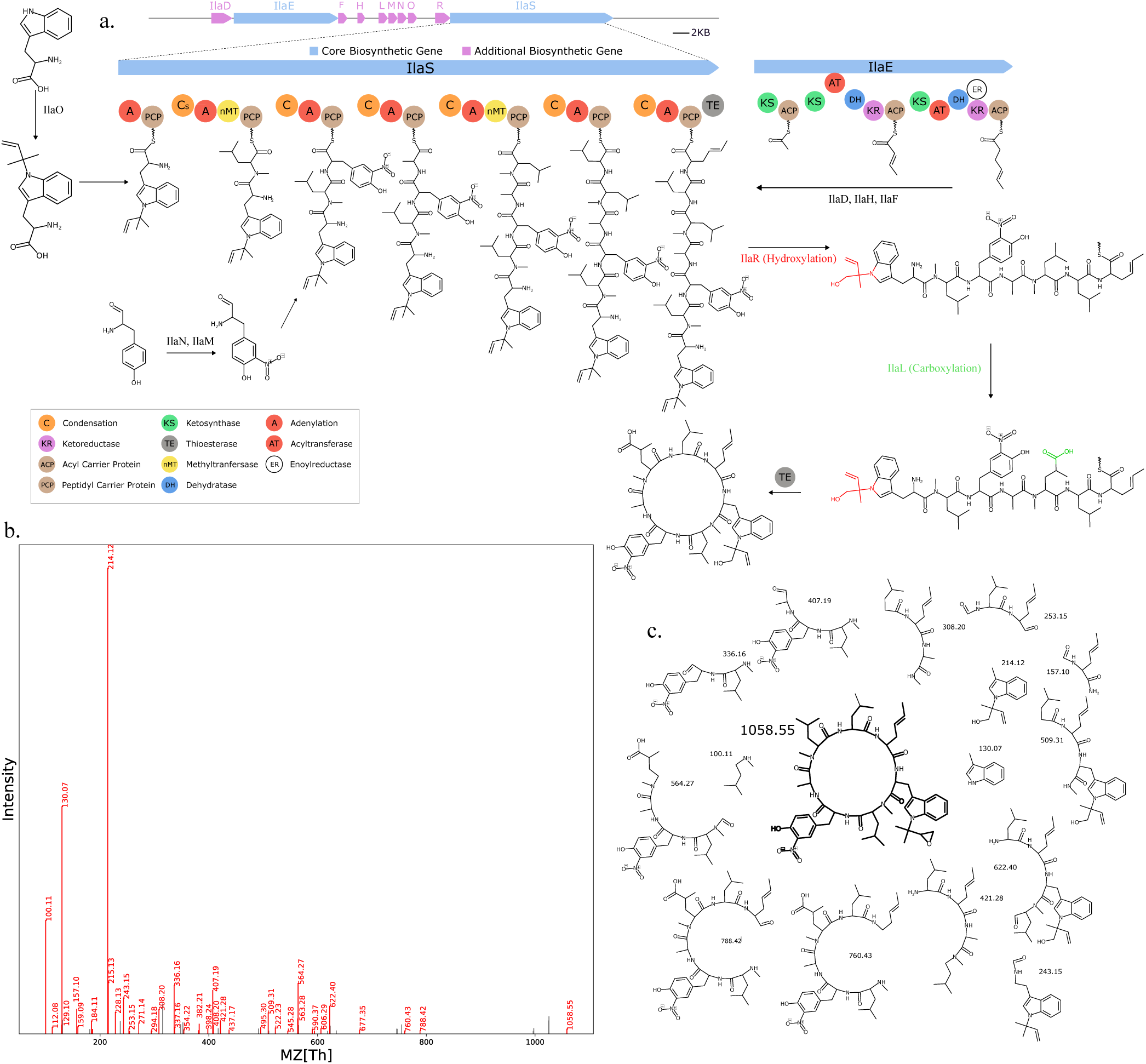
Seq2Hybrid recovers Ilamycin G BGC. a) The predicted biosynthetic pathway for Ilamycin G. b) Paired mass spectrometry data for this molecule was obtained from *Streptomyces atratus* NBRC 3897. c) Annotated mass fragments providing evidence that this molecule, or an isomer, is present in the biological sample.

**Fig. 12.**
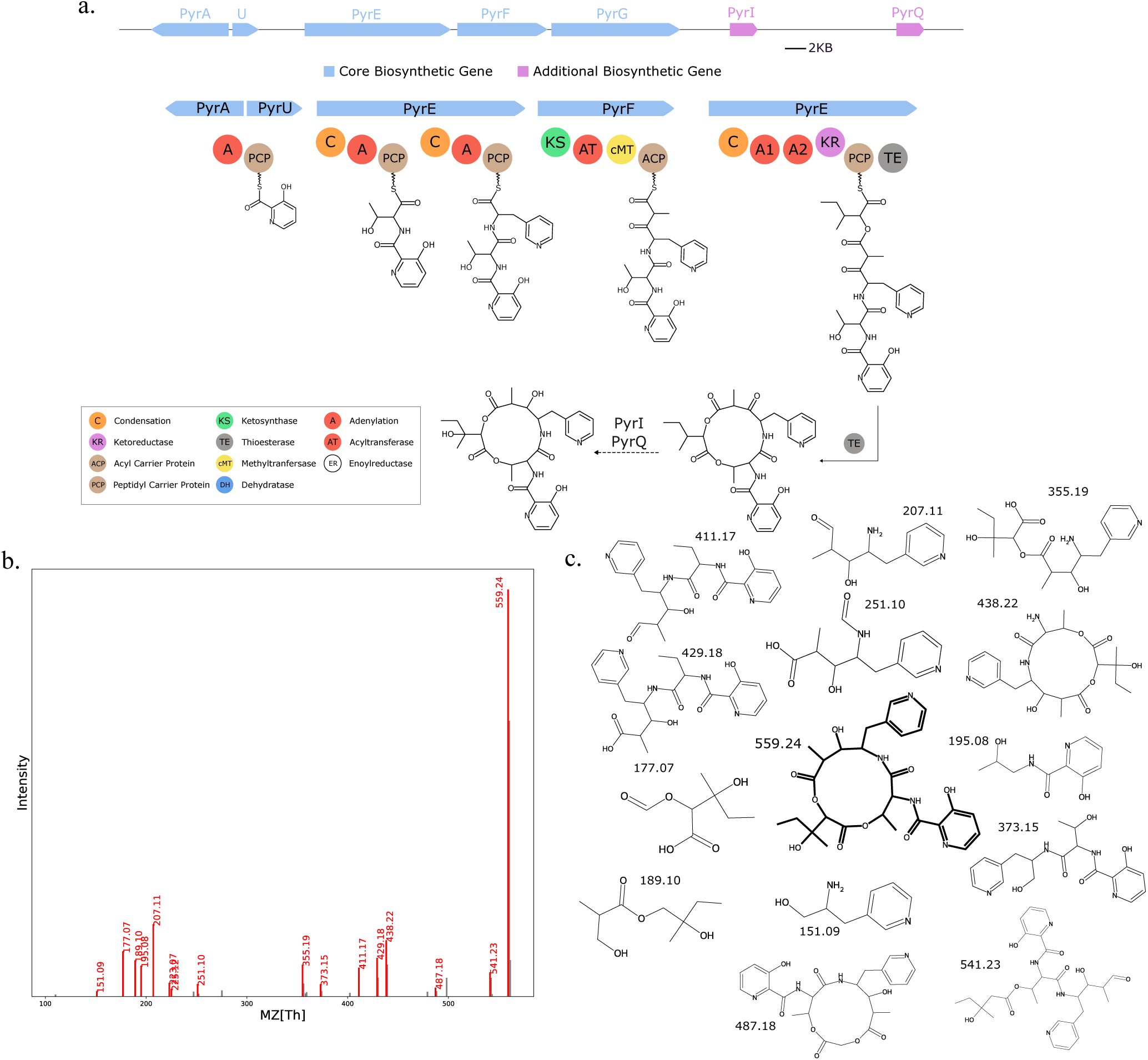
Seq2Hybrid recovers Pyridomycin BGC. a) The predicted biosynthetic pathway for Pyridomycin. b) Paired mass spectrometry data for this molecule was obtained from *Streptomyces pyridomyceticus* NRRL B-2517. c) Annotated mass fragments providing evidence that this molecule, or an isomer, is present in the biological sample.

**Fig. 13.**
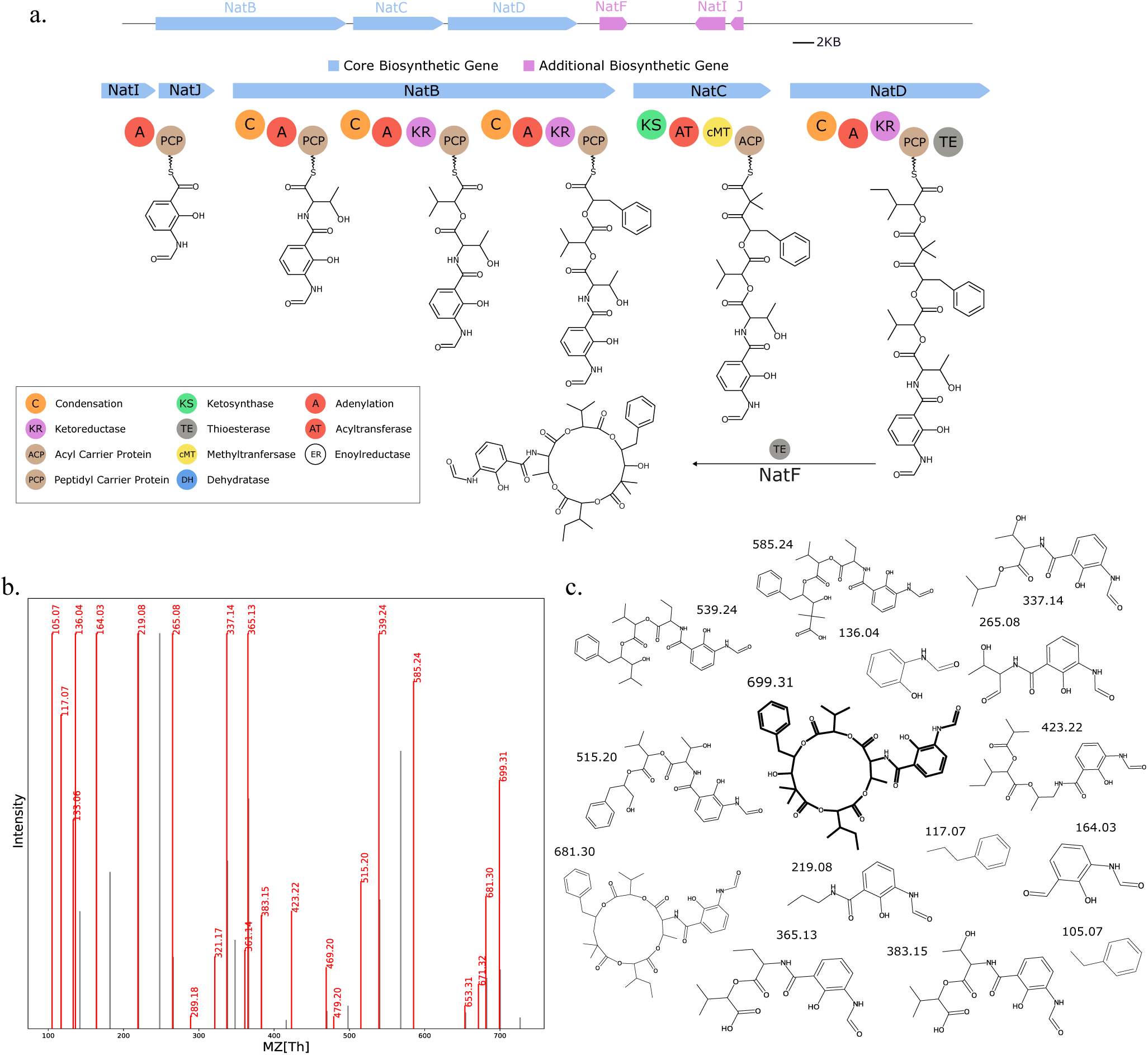
Seq2Hybrid recovers Neoantimycin BGC. a) The predicted biosynthetic pathway for Neoantimycin. b) Paired mass spectrometry data for this molecule was obtained from *Streptomyces orinoci* NBRC 13466. c) Annotated mass fragments providing evidence that this molecule, or an isomer, is present in the biological sample.

**Fig. 14.**
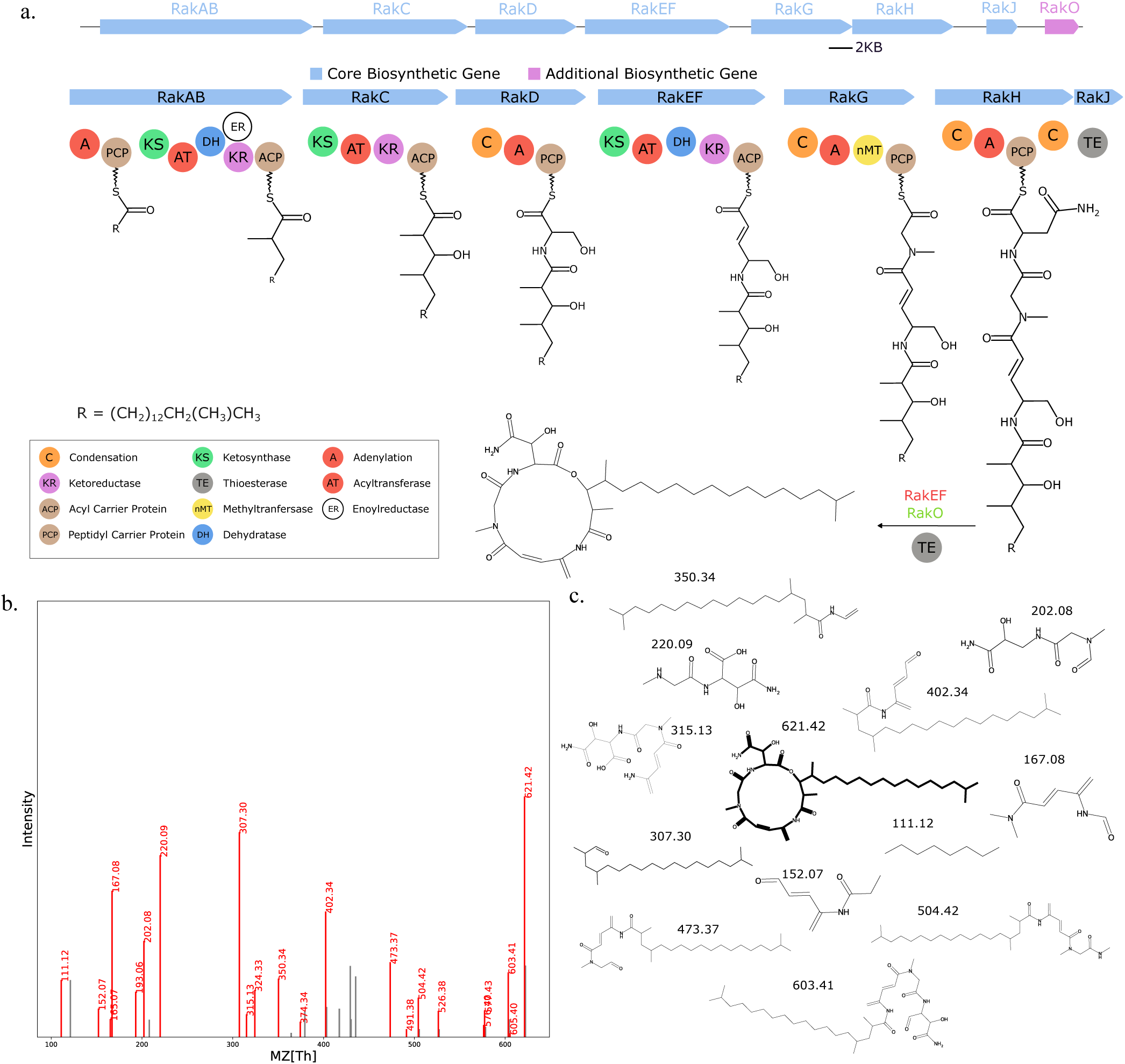
Seq2Hybrid recovers Rakicidin B BGC. a) The predicted biosynthetic pathway for Rakicidin B. b) Paired mass spectrometry data for this molecule was obtained from *Micromonospora chalcea* NRRL B-2672. c) Annotated mass fragments providing evidence that this molecule, or an isomer, is present in the biological sample.

**Fig. 15.**
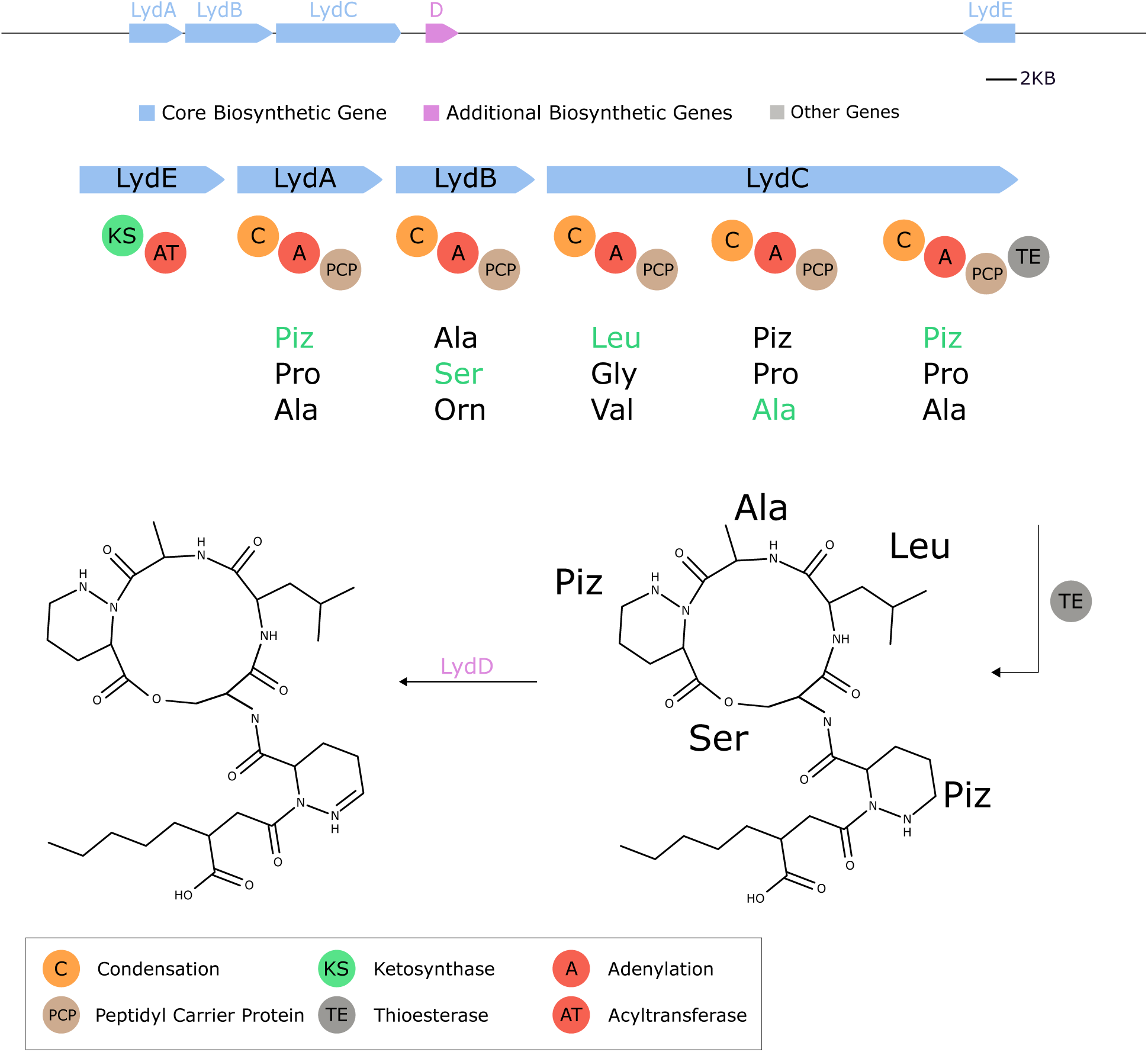
Seq2Hybrid predicts Lydiamycin BGC. The predicted biosynthetic pathway for Lydiamycin. *LydE* contains PKS-specific domains and is likely responsible for the attachment of the fatty acid tail at the N-terminus. *LydD* is a cytochrome p450 enzyme that is likely responsible for the oxidation of piperazic acid (Piz) to 2,3,4,5-tetrahydropyridazine-3-carboxylic acid.

### Seq2Hybrid identifies novel BGCs of known hybrid molecules

Seq2Hybrid identified putative BGCs for dityromycin (Fig. 7), an orphan cyclic antibiotic (58), and octaminomycin A (Fig. 16), an orphan NRP-PK hybrid with reported anti-angiogenesis effects (59), from *Streptomyces kasugaensis* NBRC 13851 and *Streptomyces hygroscopicus* NRRL B-1477 respectively. Seq2Hybrid also identified a putative BGC of origin for the immunosuppressant SW-163B, from *Streptomyces orinoci* NBRC 13466 (Fig. 17), and a putative BGC for JBIR-39 (60) in *Streptomyces violascens* NRRL B-2700 (Fig. 18).

**Fig. 16.**
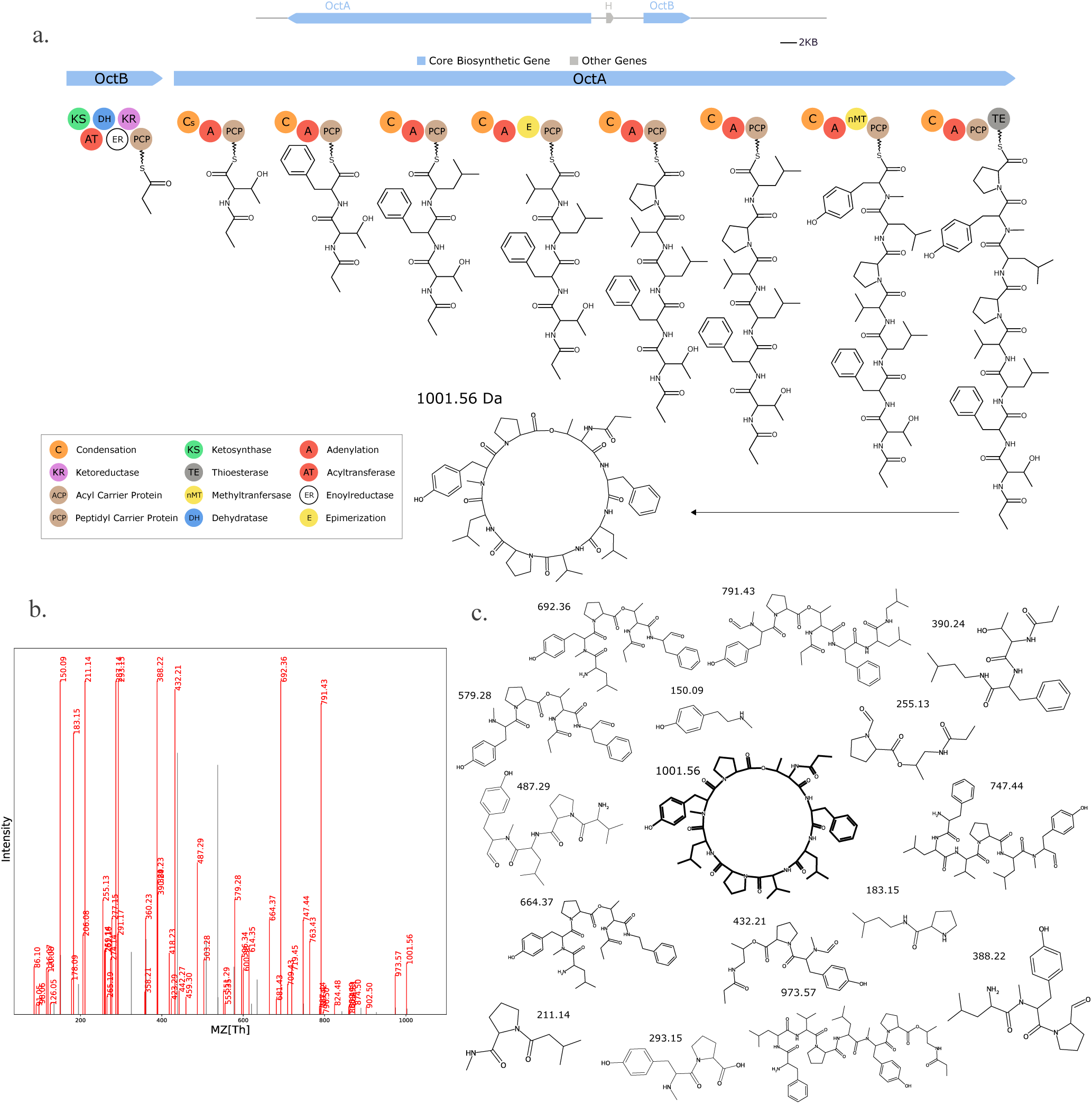
Seq2Hybrid predicts Octaminomycin A BGC. a) The predicted biosynthetic pathway for Octaminomycin A. For each residue, the top MASPR prediction matches the expected residue in Octaminomycin A. b) Paired mass spectrometry data for this molecule was obtained from *Streptomyces hygroscopicus* NRRL B-1477. c) Annotated mass fragments providing evidence that this molecule, or an isomer, is present in the biological sample.

**Fig. 17.**
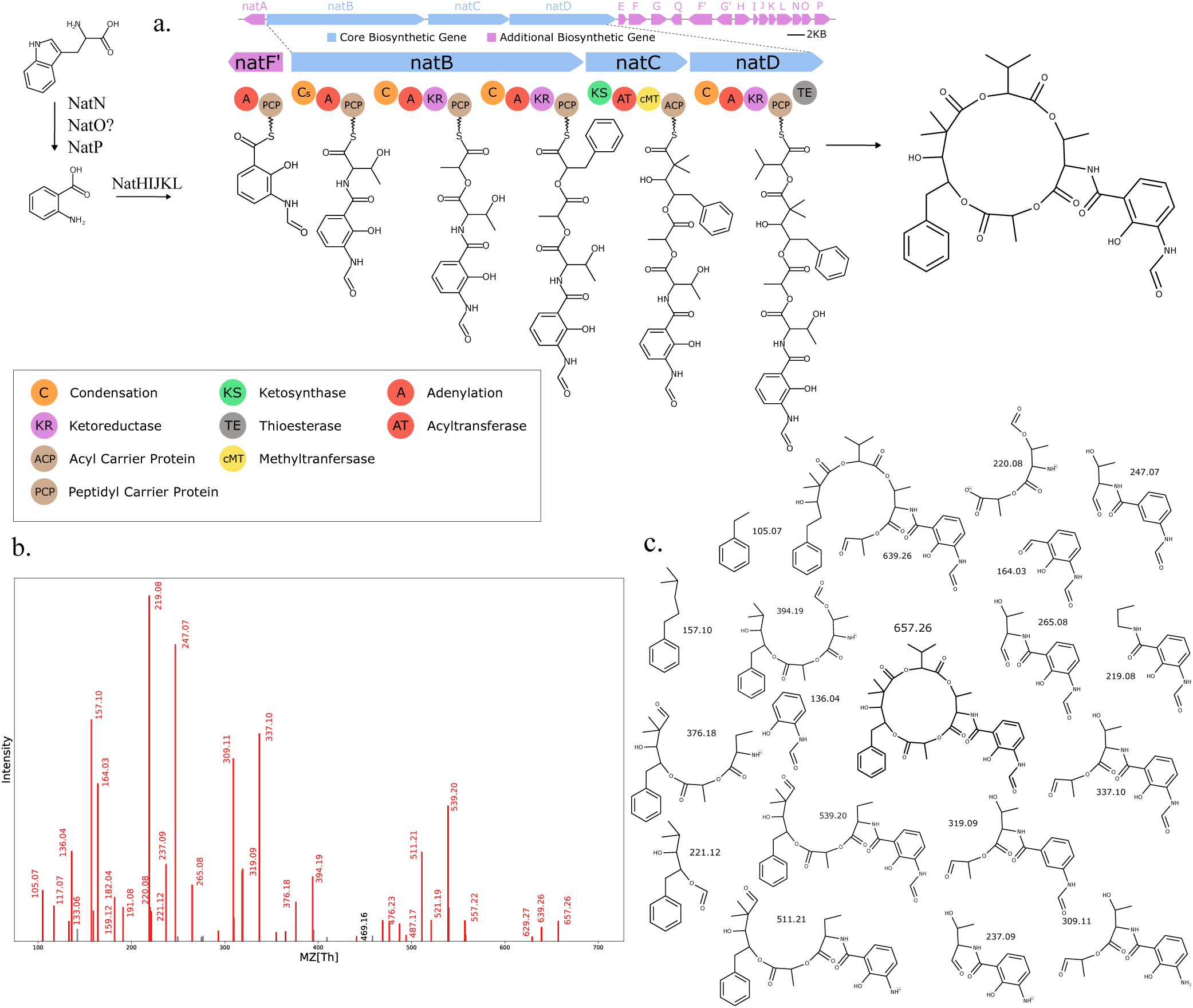
Seq2Hybrid predicts SW-163B BGC. a) The predicted biosynthetic pathway for SW-163B. b) Paired mass spectrometry data for this molecule was obtained from *Streptomyces orinoci* NBRC 13466. c) Annotated mass fragments providing evidence that this molecule, or an isomer, is present in the biological sample.

**Fig. 18.**
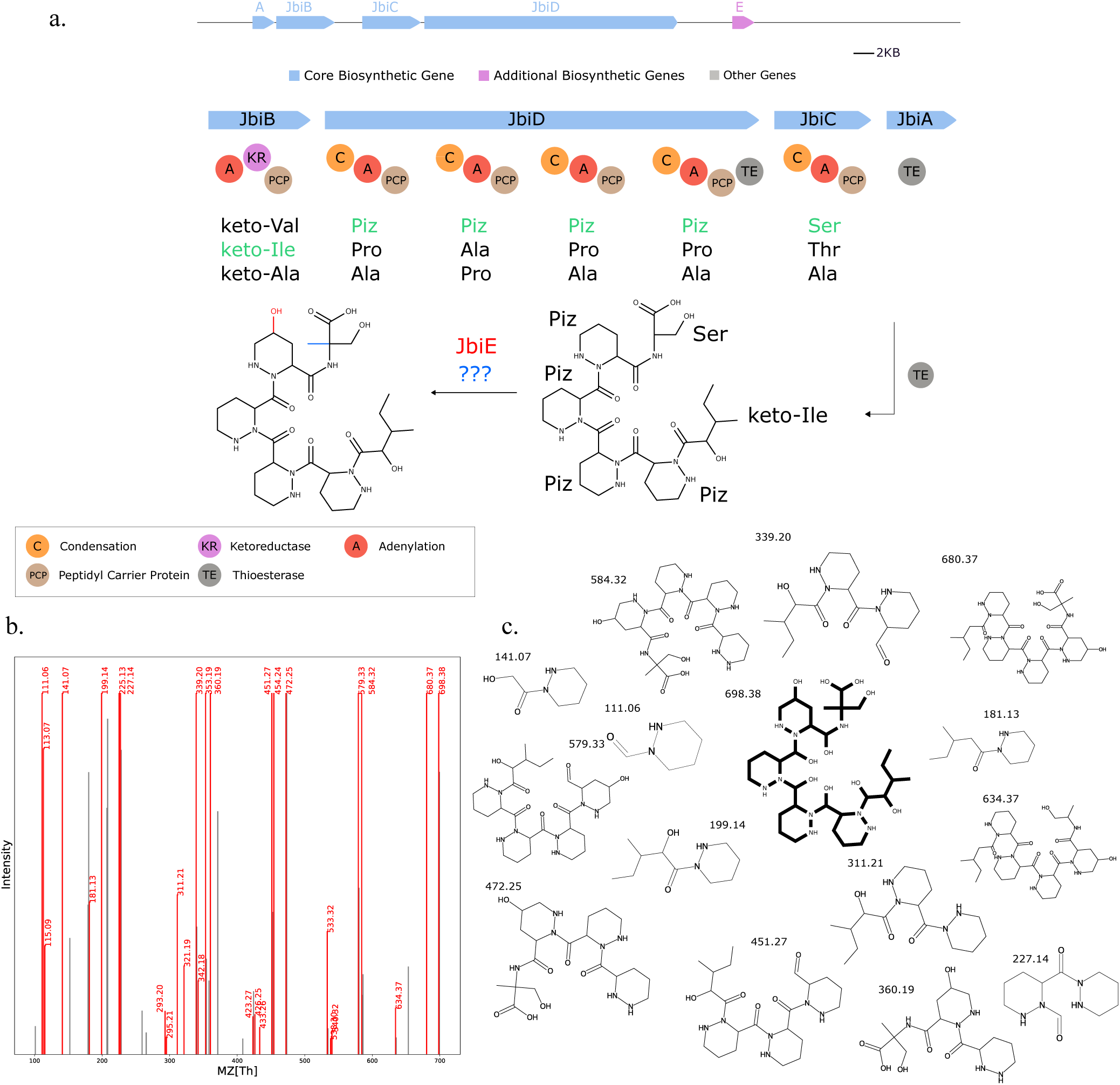
Seq2Hybrid predicts JBIR-39 BGC. a) The predicted biosynthetic pathway for JBIR-39. *JbiE* contains PKS-specific domains and is likely responsible for the attachment of the fatty acid tail at the N-terminus. Although the predicted biosynthetic pathway does not explain the C-methylation of the terminal Serine or the extra Thioesterase domain, identifying this BGC showcases the potential of MASPR for genome mining. b) Paired mass spectrometry data for this molecule was obtained from *Streptomyces violascens* NRRL B-2700. c) Annotated mass fragments providing evidence that this molecule, or an isomer, is present in the biological sample.

## Discussion

Natural products represent a goldmine of potential bioactive compounds and drug leads. Given the costly and time-consuming nature of bioactivity-based natural product discovery, *in silico* genome mining approaches are needed to fully elucidate structures encoded in hundreds of thousands of cryptic BGCs. However, existing methods that predict NRP and NRP-PK hybrid structures are affected by inaccuracies in A-domain specificity prediction and post-assembly modifications. In this work, we present MASPR, an interpretable A-domain substrate specificity predictor that achieves state-of-the-art accuracy and generalization.

By reformulating substrate classification as a regression task to predict fingerprints, MASPR improves on the accuracy of existing methods by up to 15%. Since MASPR is trained to generate A-domain-specific substrate embeddings from molecular fingerprints which are computable from SMILES representations, it can create substrate embedding databases that include substrates not present in the training data. This enables MASPR to perform zero-shot prediction of novel substrates by computing an embedding for a given A-domain and searching for the nearest substrates in the embedding database. In a leave-one-substrate-out cross-validation study designed to benchmark zero-shot performance, MASPR achieved higher than 50% top-5 accuracy for over half of the held-out substrates. Because MASPR is trained to predict substructure-based molecular fingerprints, its predictions are interpretable, as we can annotate substructures in a given substrate that are the most relevant in prediction. MASPR performance significantly drops when using a one-hot featurization, showing that the per-residue ESM-2 embeddings encode signal relevant to A-domain specificity. Our results further indicate that averaging per-residue embeddings across the entire A-domain can dampen this signal, and the highest accuracy is achieved when maintaining a separate channel for each Stachelhaus residue.

We used MASPR to develop Seq2Hybrid, a genome mining pipeline for discovering novel NRP-PK hybrid structures encoded in microbial genomes. NRP-PK hybrids are a class of natural products with promising therapeutic value, yet they remain underrepresented in genome mining approaches due to their complex biosynthesis and structure. Seq2Hybrid recovered known hybrid molecules from BGCs with much higher accuracy than existing approaches and connected orphan compounds octaminomycin A, dityromycin, JBIR-39, and SW-163B to their respective BGCs of origin. Seq2Hybrid predictions are further filtered with paired mass spectrometry data and error corrected using variable mass spectral database search methods.

At present, MASPR’s substrate specificity predictions are based solely on A-domain sequences. Future enhancements could incorporate NRP-specific biosynthetic logic into the prediction model. For example, an A-domain followed by a KR-domain is likely to recruit a keto-acid, and an A-domain preceded by a heterocyclization domain is more likely to incorporate Serine, Threonine, or Cysteine. Integrating three-dimensional substrate information is another potential avenue for improvement, as the current training data and fingerprint encodings cannot differentiate between stereoisomers. Finally, Seq2Hybrid is currently limited to predicting modular NRP-PK hybrids, and it cannot process the iterative synthesis often observed in type 2 PK hybrids. Addressing this limitation would make MASPR and Seq2Hybrid applicable to a wider variety of microbial BGCs.

## Code Availability

Training data and code for MASPR are available here: https://github.com/abhinadduri/MASPR. MASPR is also available on the web via Colab notebook: https://bit.ly/colab-maspr.

Seq2Hybrid is supported as a web service at https://run.npanalysis.org. Train and test BGCs and their corresponding Seq2Hybrid predictions are available as supplementary files and on Google Drive: https://bit.ly/seq2hybrid-bgc-predictions.

## Methods and Supplementary Info

### Curating training data for MASPR

Training data for MASPR was obtained from MIBiG 3.0, which contains substrate specificity annotations for A-domains. For promiscuous A-domains that recruit multiple monomers, each pair between the A-domain and monomers was treated as a training data point, resulting in 2294 data points. For each A-domain, we used the ESM-2 model `esm2_t33_650M_UR50D` (53) to extract 1280-dimensional per-residue embeddings from the entire sequence. Then, we performed a sequence alignment to a reference A-domain (PDBID:*1AMU*) and extracted the embeddings corresponding to the Stachelhaus residues to obtain a 10 *×* 1280 embedding for each A-domain sequence in the training data.

### Training procedure for MASPR

MASPR was implemented and trained in PyTorch 2.0 using frozen A-domain embeddings as input. To train the fingerprint predictor neural network, each substrate in the training data was converted to a 296-dimensional fingerprint representation, where the first 167 entries correspond to the MACCS key of the substrate, the next 128 entries correspond to a Morgan fingerprint with a radius of 2, and the last entry corresponds to an average partial charge of the atoms in the substrate. Although this fingerprint was chosen as the combination of RDKit descriptors that led to the best performance, clustering the substrates in the training data using *L*2-distance between fingerprints (Fig. 19) recovers previously reported A-domain-specific clustering of amino acids (22, 32).

**Fig. 19.**
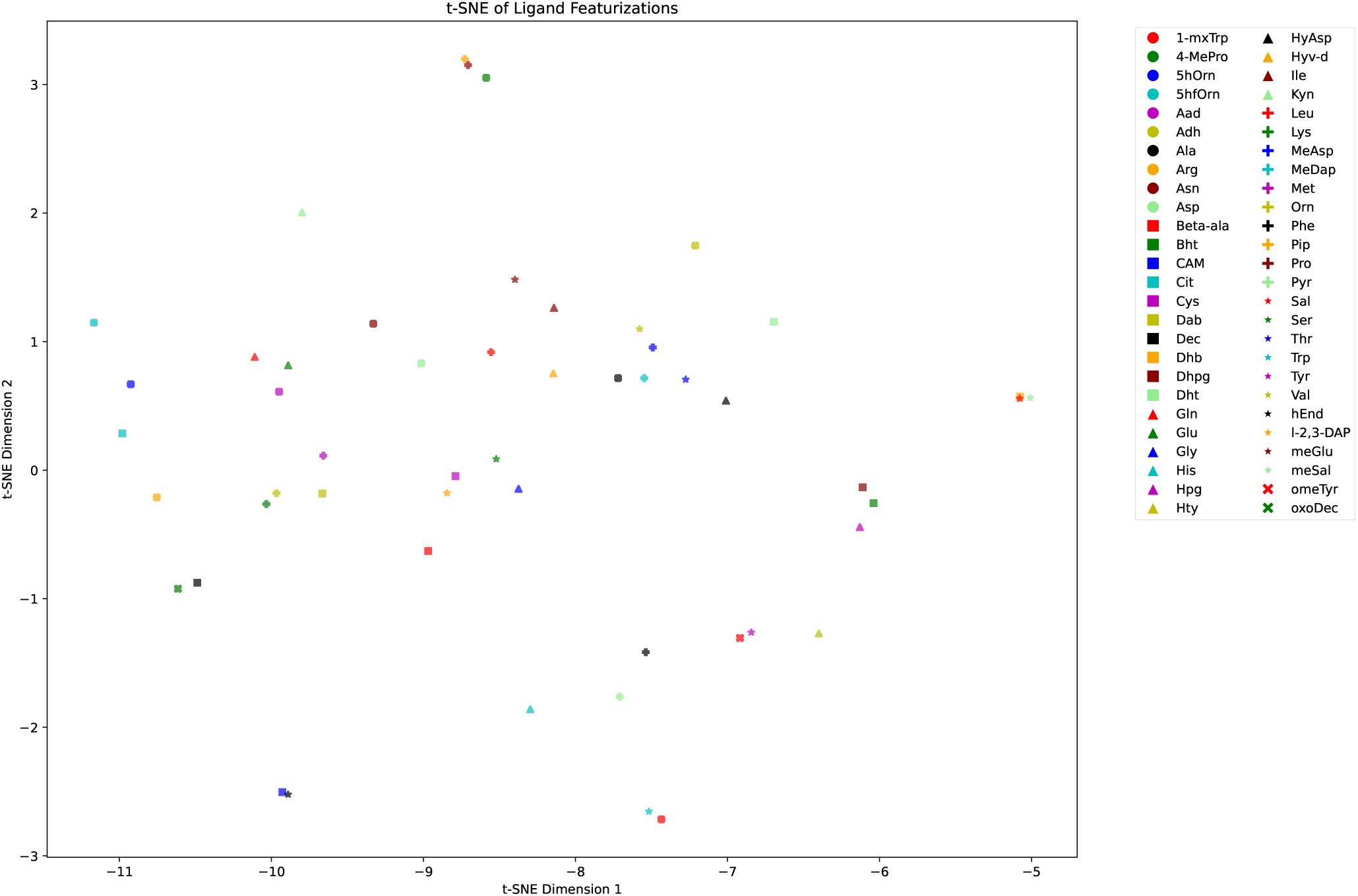
tSNE visualization of substrate fingerprints in the training data. Points in the visualization are semantically well separated and recapture previously reported A-domain-binding specific similarities across ligands (22, 32).

The fingerprint representation is used as a regression target during the training. The architecture of the neural network includes several layers: Linear (1280 *×* 480), Linear (480 *×* 240), Flatten (across the 10 Stachelhaus residues), Linear (2400 *×* 240), Linear (240 *×* 240), and Linear (240, 296). Each Linear layer, except the last, is followed by an ELU activation and a LayerNorm to facilitate training stability and performance. The network was trained for 80 epochs using cosine distance as the loss function, with Adam optimizer and weight decay (AdamW optimizer in PyTorch), a learning rate of 0.0001, and exponential decay of 0.8 every ten epochs. Cosine distance loss was implemented as `(1 - CosineSimilarity(predicted_fingerprint, target_fingerprint))`, averaged across all bits of the fingerprint.

The classifier head neural network is trained concurrently with the fingerprint predictor network. Each forward pass of the fingerprint predictor is followed by the classifier head being trained to predict the substrate label from both the predicted and target fingerprints, meaning the classifier network parameters are updated twice during each forward pass of the fingerprint predictor network. The classifier head’s architecture comprises two linear layers: the first maps 296 inputs to 296 outputs. This is followed by a ReLU activation function, whose output is used as the A-domain-specific embedding for the nearest substrate search. The second linear layer maps 296 to 41 outputs, corresponding to the number of unique substrates in the data. The model is trained with CrossEntropyLoss with a learning rate of 0.0001. The weights for the first linear layer are initialized to the 296 *×* 296 identity matrix. For benchmark models that replaced the nearest substrate search by directly classifying a substrate label, the classifier head network is omitted, and the final output dimension for the fingerprint predictor network is changed from 296 to 41. We used a batch size of 128 for all models.

To perform top-*k* substrate specificity classification for a given A-domain, MASPR first computes the predicted embedding using the fingerprint predictor network. Then, MASPR feeds the predicted fingerprint into the classifier head and extracts the hidden layer embedding. The hidden embedding is compared against a database of substrate embeddings and the *k* closest substrates are returned, where the distance between substrates is computed as the cosine distance between their embeddings.

### NRP-PK Hybrid BGC detection

Seq2Hybrid identifies BGCs by searching for both NRP-specific and PKS-specific domains. The NRP-specific domains include adenylation (A-) domains, which are responsible for incorporating either an amino acid or hydroxy acid into the growing natural product, condensation (C-) domains, which are involved in peptide bond formation, and peptidyl carrier protein (PCP-) domains, which are responsible for transporting the intermediate natural product to downstream domains. The PKS-specific domains include the acyltransferase (AT-) domain, which recruits an *α*-carboxyacyl-CoA extender unit (or ketide unit), the acyl carrier protein (ACP-) domain, which accepts the ketide unit from the AT-domain, and the ketoacyl synthase (KS-) domain, which catalyzes the carbon-carbon bond between the growing natural product and the new ketide-ACP intermediate (61). We created a database of 278 A-, C-, and PCP-domains and 183 AT-, KS-, and ACP-domains stored as profile HMMs (62). Seq2Hybrid searches for these domains in six-frame translations of each contig in the input genome. Each identified domain is extended upstream and downstream by a user-specified threshold (10KB by default), and the overlapping genome segments are merged. Seq2Hybrid reports the resulting segments as the BGC regions.

### Annotating active domains with monomer specificity

In NRP-PK hybrid synthesis, A-domains incorporate amino acids or hydroxy acids and AT-domains introduce ketide units into the growing molecule. Seq2Hybrid predicts the monomers that are most likely to be recruited by each active domain in the hybrid BGC. Since A-domains can be promiscuous in their substrate selection (22), we annotate each A-domain with its top three MASPR predictions. To predict AT-domain specificity, Seq2Hybrid uses a strategy similar to Minowa et al. (17) to extract a 24 amino acid signature of the active site of AT-domains. Seq2Hybrid forms a single HMM profile with all reference AT-domains, and extracts the signature via alignment against this HMM profile. Then, Seq2Hybrid uses a random forest to predict substrate specificity given the one-hot encoded signature of the AT-domain.

### Biosynthetic gene graph

Although certain genes may be inactive in the biosynthetic pathway of NRPs, the sequential arrangement of amino acids in the NRP matches the order in which they appear in the BGC (i.e. co-linearity) (63). Taking *n* as the total number of biosynthetic genes, and allowing for the possibility of *k* inactive genes, this yields an upper limit of 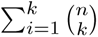 potential gene assembly orders. This number is higher in PKs, typically due to non-linearity in gene-to-gene interactions. With a total of *n* genes and up to *k* inactive genes, up to 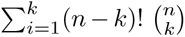 assemblies are possible for PKs. This number escalates rapidly for large values of *n*. To address this, Seq2Hybrid uses a *biosynthetic gene graph*, constructed on the genes within the BGC. In this graph, nodes represent genes, and an edge between two nodes *s* and *t* signifies that gene *t* can follow gene *s* in the final biosynthesis.

First, Seq2Hybrid adds an edge from *s* to *t* if gene *s* ends in a C-terminal communication-mediating (COM) domain and gene *t* begins with an N-terminal COM domain, as these domains enable gene-to-gene interactions (64, 65) (Rule 1). Second, Seq2Hybrid adds an edge from *s* to *t* if gene *s* ends with a C-domain (A-domain) and *t* starts with an A-domain (PCP-domain) (Rule 2). Third, Seq2hybrid adds an edge from *s* to *t* if gene *t* is downstream of *s* on the same strand (Rule 3). Fourth, Seq2Hybrid adds an edge from any gene *s* to gene *t* if *t* contains a thioesterase or thioreductase domain (Rule 4). Fifth, if gene *s* is a singleton domain (i.e. contains only a single active domain), then Seq2hybrid adds an edge from *s* to all genes *t* (Rule 5). Finally, Seq2Hybrid trims the graph by removing outgoing edges starting from release domains. Furthermore, if a gene *t* starts with an N-terminal COM domain and a gene *s* does not end with a C-terminal COM domain, then Seq2Hybrid removes the edge from *s* to *t* unless this would leave *t* with no incoming edges (Fig. 20).

**Fig. 20.**
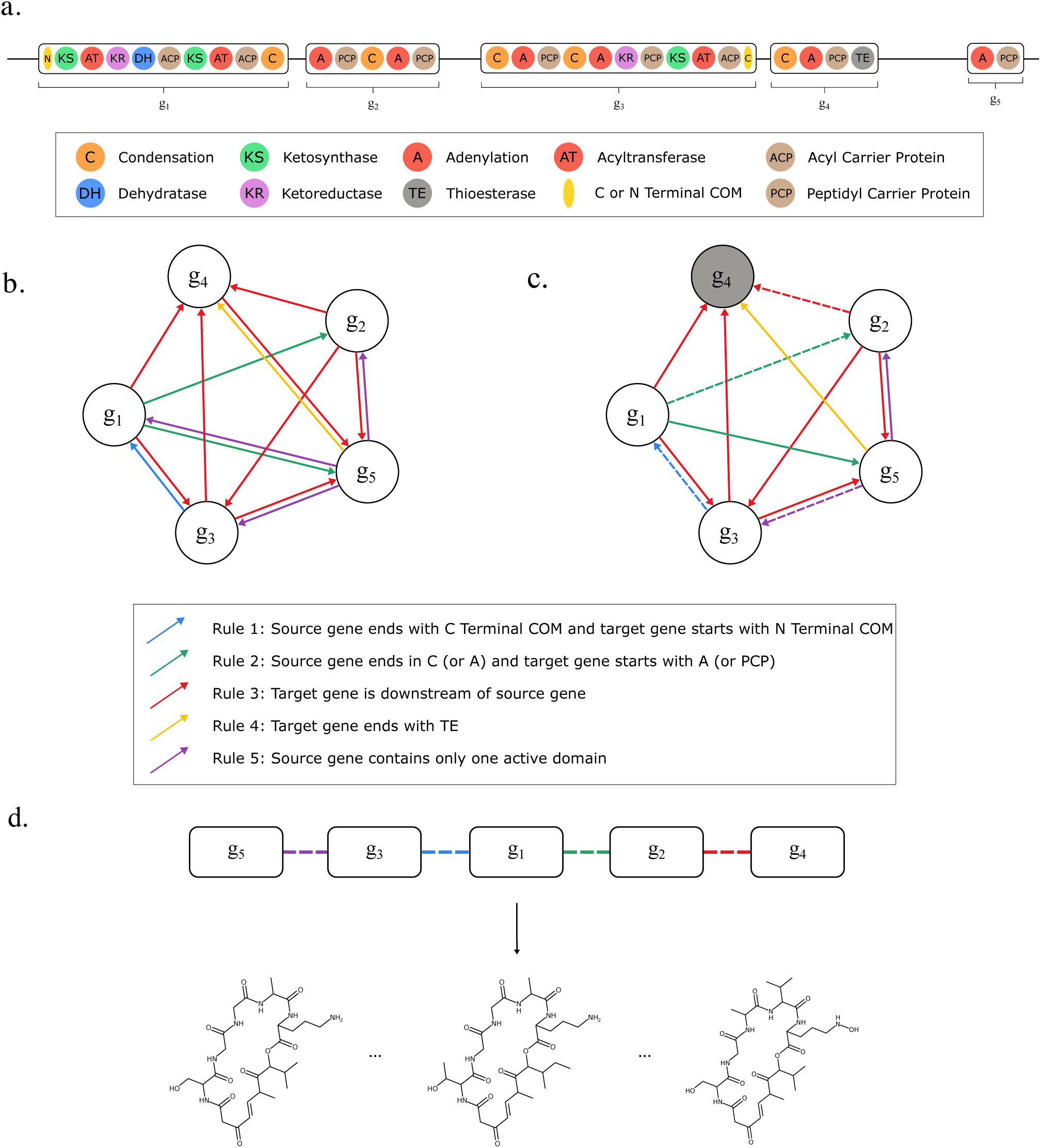
Constructing biosynthetic gene graphs. Starting from the BGC (a), the biosynthetic gene graph (b) is constructed using five rules. (c) The biosynthetic gene graph is then trimmed. The edge from *g*_1_ to *g*_5_ is removed since *g*_4_ ends with a TE domain. The edge from *g*_5_ to *g*_1_ is removed since *g*_1_ starts with an N-terminal COM domain, but *g*_5_ does not end with a C-terminal COM domain. The sink node is highlighted in grey. One of the feasible assembly orders is depicted with dashed lines. d) Candidate assembly line orders are extracted from the biosynthetic gene graph.

### Traversing the biosynthetic gene graph

Upon constructing the biosynthetic gene graph, Seq2Hybrid performs traversals through the graph to recover different biosynthetic assemblies. First, Seq2Hybrid identifies a sink node within the graph, which is defined as the node with the minimal number of outgoing edges (Fig. 20*c*). Let *n* denote the total number of nodes in the graph. Since certain genes can be inactive in the final biosynthesis (22), let *k* denote the user-specified number of allowed inactive genes. By default, *k* is set to the count of singleton domains within the BGC. Then, Seq2Hybrid reports all paths in the graph that terminate at this sink node and contain at least *n− k* nodes. The resulting paths are referred to as *assembly lines*.

### Generating hybrid cores from assembly lines

Given an assembly line, Seq2Hybrid considers various assignments of monomers to each active domain to yield final molecular structures. Seq2Hybrid considers the top three predictions for each A-domain and the top one prediction for each AT-domain. With *n* A-domains in an assembly line, this results in 3*^n^* different monomer assignments per assembly line. To limit the computational complexity for large values of *n*, Seq2Hybrid adopts a dynamic programming scheme (22) to only consider the top *s* highest scoring monomer assignments (by default *s* = 1000). The score of a monomer assignment is calculated as the sum of the scores of individual monomers at each A-domain in the assembly, predicted by AdenPredictor. In practice, *n* is typically smaller than eight, making it feasible to consider all combinations. For each unique monomer assignment, we construct a *core molecule* by connecting the respective monomers chosen in that assignment (Fig. 21).

**Fig. 21.**
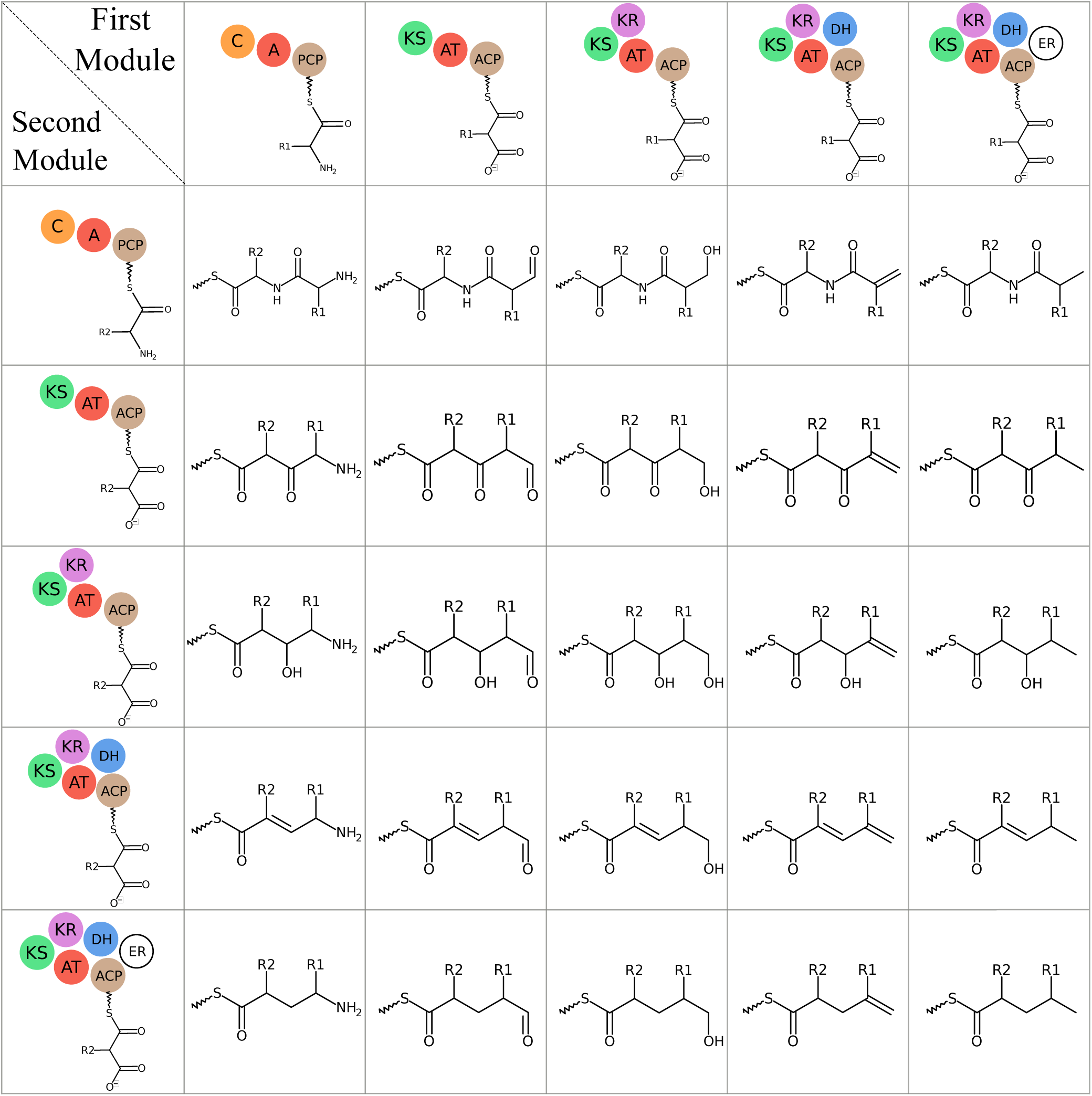
Connections between different types of monomers in hybrid assembly. The connections formed between amino acids and ketides that have been successively reduced. Hydroxy acids are connected to other monomers in the same way as amino acids, with the exception that the nitrogen is replaced by oxygen.

After the construction of the hybrid cores, various modifications are applied based on the presence of modification domains. These modifications include formylation (via N-terminal F-domain), N-methylation (via methylation domain after A-domain), C-methylation or O-methylation (via methylation domain after AT-domain), thiazoline or oxazoline ring formation (via heterocyclization domain after A-domain) (66), further ring oxidation or reduction (via oxidation domain or reduction domain after heterocyclization domain), and successive reductions of ketides to hydroxyl, alkene, and methylene groups (via ketoreductase, dehydratase, and enoylreductase domains, respectfully) (67).

### Applying pre-assembly modifications

Certain non-standard amino acids are synthesized via enzymes or other biosynthetic genes present in the BGC. For example, the biosynthesis of Ilamycins G (Fig. 11) relies on the production of 2-aminohex-4-enoic acid from a set of PK biosynthetic genes in the same BGC. Such modifications of monomers on the assembly line are referred to as pre-assembly modifications. Seq2Hybrid considers monomers with known pre-assembly modifications if the specific enzymes required for those modifications are present in the BGC. Seq2Hybrid accounts for 41 monomers collected using a literature review (Supplementary Table S3), and uses the zero-shot capability of MASPR to predict specificity outside of these monomers.

### Applying post-assembly modifications

After the assembly of core hybrid molecules, enzymes present in the BGC apply various post-assembly modifications to the core molecular structures. To account for common modifications in hybrid biosynthesis, Seq2Hybrid accounts for 170 NRP-specific modifications, 93 PKS-specific modifications, and 41 NRP-PK-specific modifications mined from the natural product literature (Fig. 22). Seq2Hybrid also collects information on the enzymes that perform these modifications and stores them as profile HMMs. For a given hybrid BGC, Seq2NRP only considers a modification if all of its required enzymes are present in the BGC. This dramatically reduces the number of required modifications to consider when predicting the structure of the mature natural product. For each considered modification, Seq2Hybrid uses the Ullman algorithm (68) to compute a subgraph isomorphism between the core and the motif of the modification. This is computationally tractable as the graph sizes for the cores and motifs are small. To compute the set of mature natural products, Seq2Hybrid combinatorially applies non-overlapping modifications to each core molecule (Fig. 23).

**Fig. 22.**
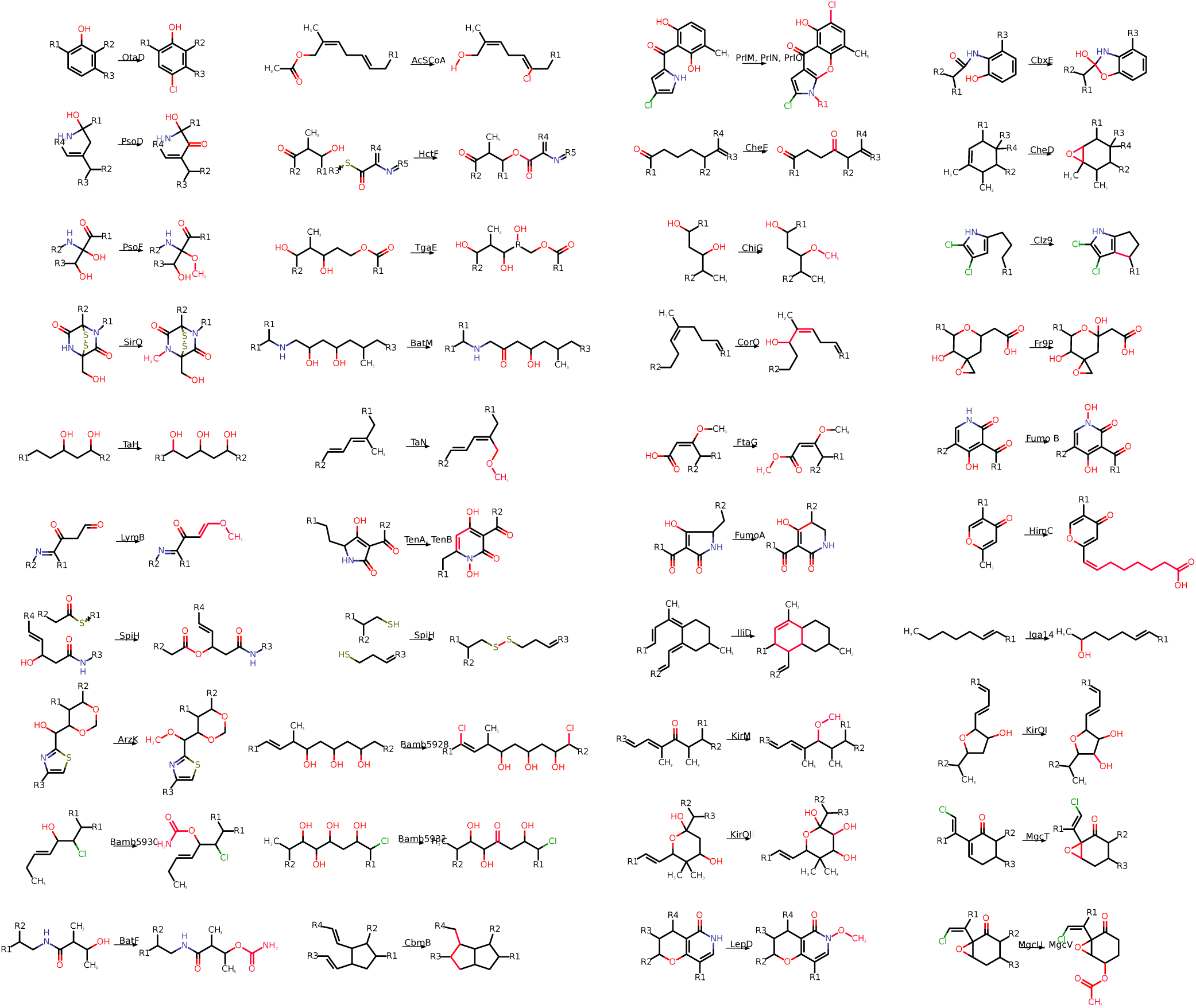
A subset of the modifications collected from the literature. Modifications were collected by searching the literature for common enzymatic reactions in NRP, PK, and NRP-PK hybrid natural product synthesis.

**Fig. 23.**
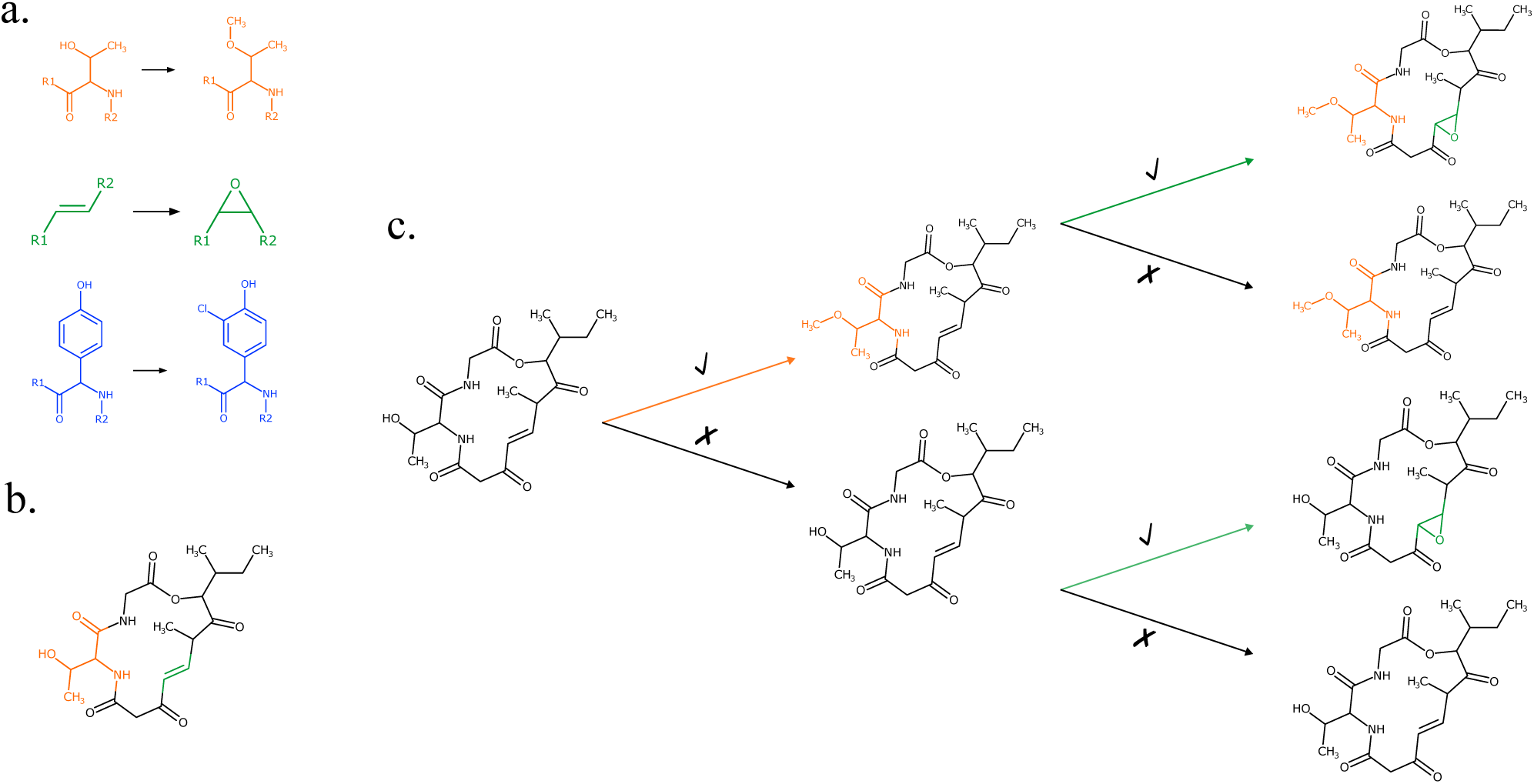
Post-assembly modification of core molecules to mature NRP-PK hybrids. (a) Starting with an initial set of three modifications, (b) modifications are mapped to a core molecule via subgraph isomorphism. The blue modification is discarded as it does not map to the core. The orange and green modifications map to a single site in the molecule. (c) Seq2Hybrid combinatorically applies the modifications that are not overlapping.

---

**Supplementary Table S1.**
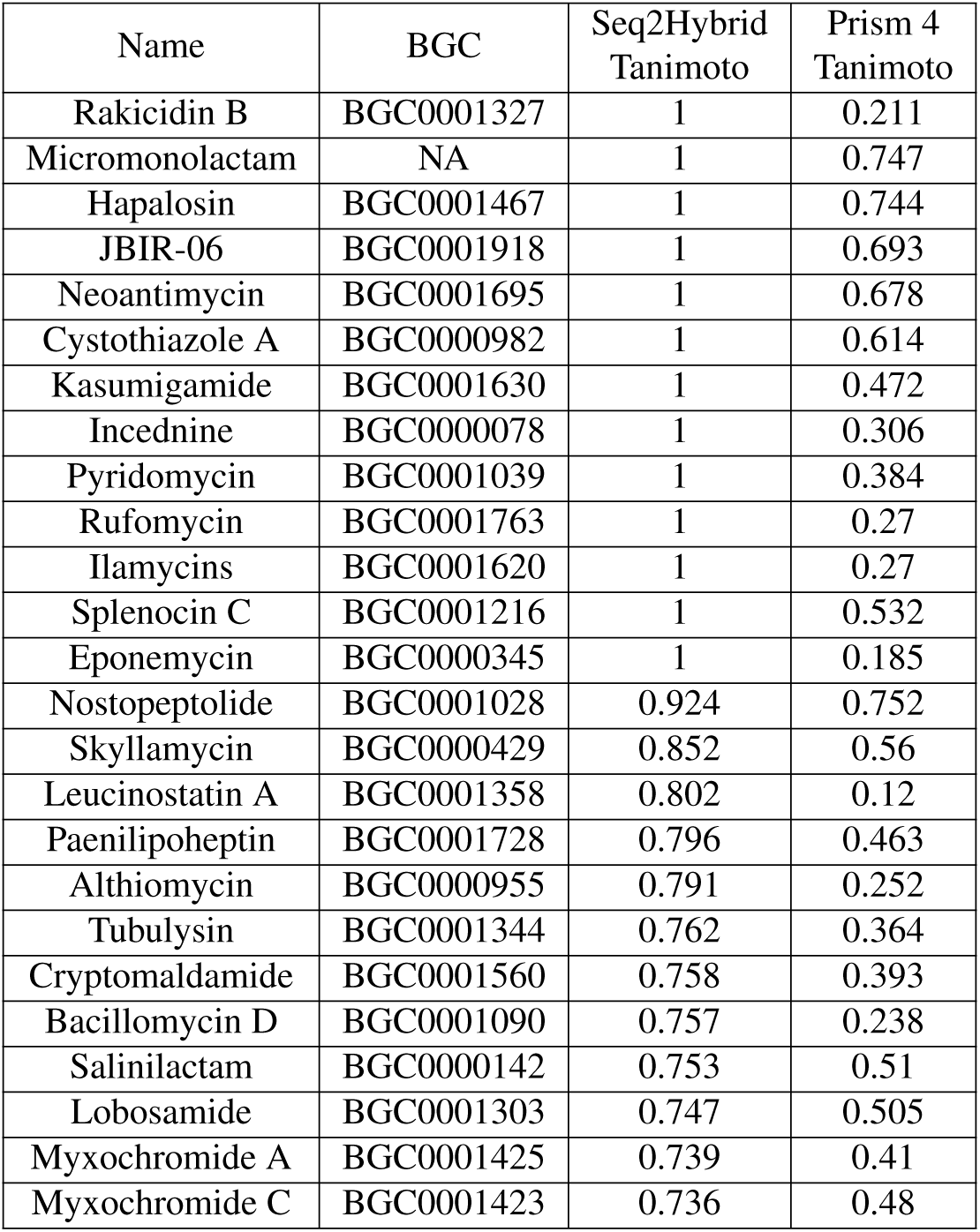
Seq2Hybrid vs PRISM 4 performance at recovering NRP-PK hybrid molecules at or above a Tanimoto similarity threshold of 0.7 in either the training or test set.

**Supplementary Table S2.**
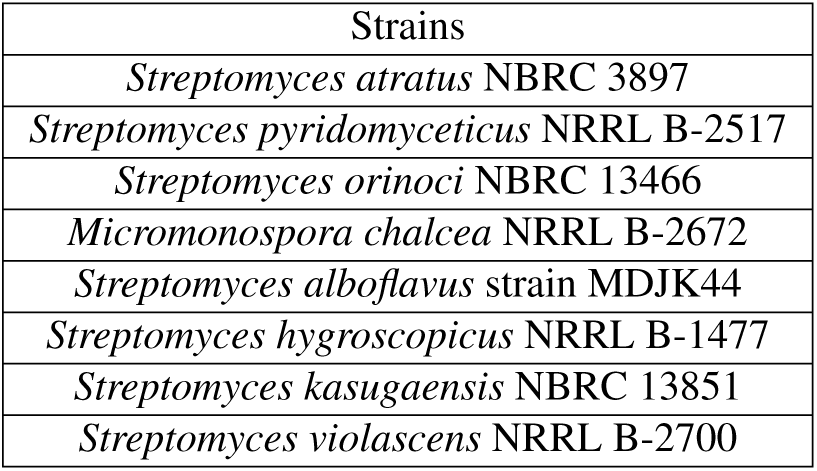
Strains used in this study for which mass spectrometry data was obtained.

**Supplementary Table S3.**
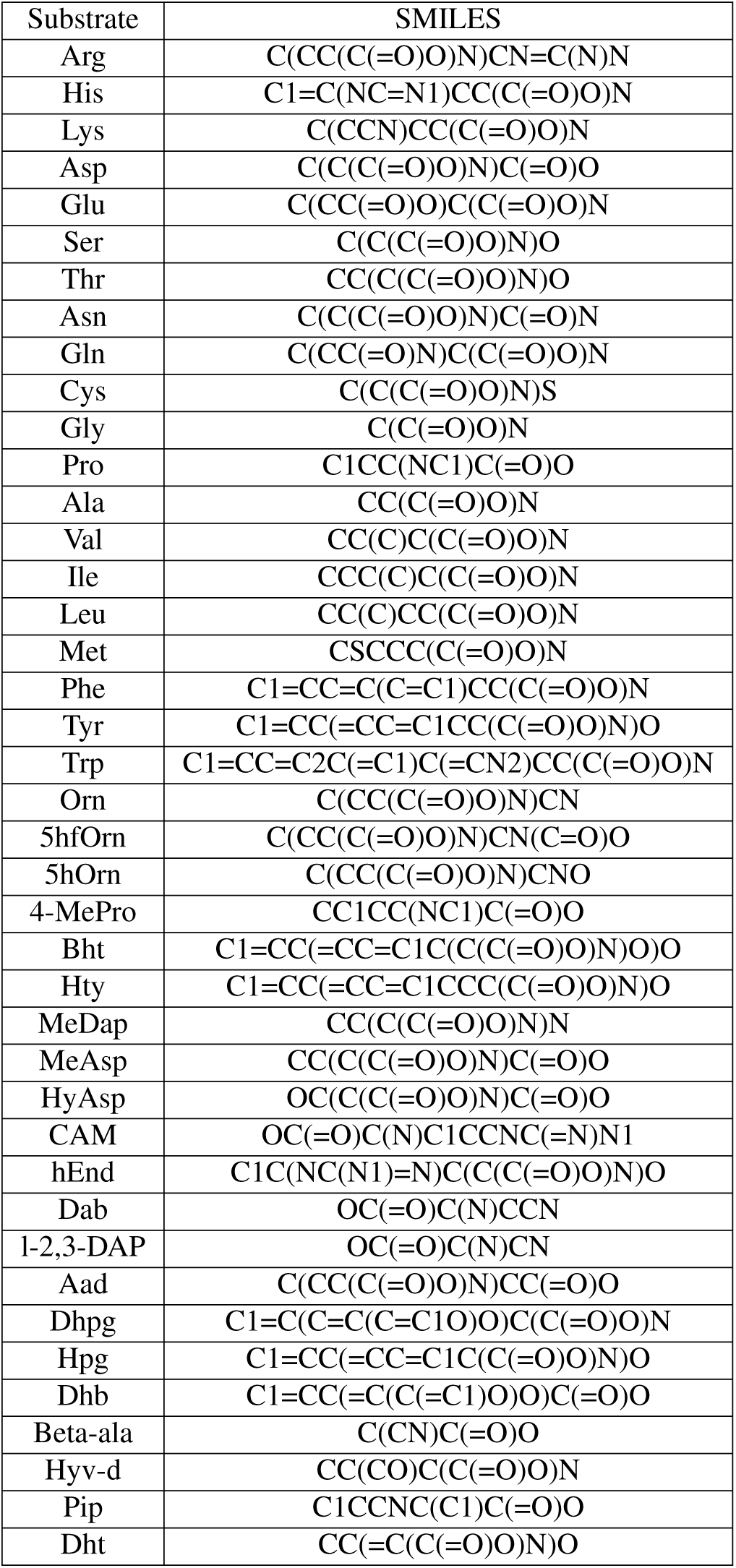
Substrates used in this study to train MASPR and their corresponding SMILES representations.

